# A *qnr*-plasmid allows aminoglycosides to induce SOS in *Escherichia coli*

**DOI:** 10.1101/2021.04.29.441945

**Authors:** Anamaria Babosan, David Skurnik, Anaëlle Muggeo, Gerald B. Pier, Zeynep Baharoglu, Thomas Jové, Marie-Cécile Ploy, Sophie Griveau, Fethi Bedioui, Sébastien Vergnolle, Sophie Moussalih, Christophe de Champs, Didier Mazel, Thomas Guillard

**Affiliations:** Inserm UMR-S 1250 P3Cell, SFR CAP-Santé, Université de Reims-Champagne-Ardenne, Reims, France; Institut Necker-Enfants Malades, Inserm U1151-Equipe 11, Université Paris Descartes, Paris, France. Laboratoire de Bactériologie, AP-HP, Hôpital Necker-Enfants Malades, Paris, France. Division of Infectious Diseases, Department of Medicine, Brigham and Women’s Hospital, Harvard Medical School, Boston, MA, USA; Inserm UMR-S 1250 P3Cell, SFR CAP-Santé, Université de Reims-Champagne-Ardenne, Reims, France. Laboratoire de Bactériologie-Virologie-Hygiène Hospitalière-Parasitologie-Mycologie, CHU Reims, Hôpital Robert Debré, Reims, France; Division of Infectious Diseases, Department of Medicine, Brigham and Women’s Hospital, Harvard Medical School, Boston, MA, USA; Institut Pasteur, Unité Plasticité du Génome Bactérien, CNRS UMR3525, Paris, France; Université de Limoges, Inserm, CHU Limoges, RESINFIT, UMR 1092, Limoges, France; Chimie ParisTech, PSL Research University, CNRS, Institute of Chemistry for Life and Health Sciences, Paris, France; Laboratoire d’Hématologie, CH de Troyes, Troyes, France

## Abstract

The plasmid-mediated quinolone resistance (PMQR) genes have been shown to promote high-level bacterial resistance to fluoroquinolone antibiotics, potentially leading to clinical treatment failures.

In *Escherichia coli*, sub-inhibitory concentrations (sub-MIC) of the widely used fluoroquinolones are known to induce the SOS response. Interestingly, the expression of several PMQR *qnr* genes is controlled by the SOS master regulator.

During the characterization of a small *qnrD*-plasmid carried in *E. coli,* we observed that the aminoglycosides become able to induce the SOS response in this species, thus leading to the transcription of *qnrD*.

We found that induction of the SOS response is due to nitric oxide (NO) accumulation in presence of sub-MIC of aminoglycosides. We demonstrated that the NO accumulation is driven by two plasmid genes, ORF3 and ORF4, whose products act at two levels. ORF3 encode a *FAD*-binding oxidoreductase which helps NO synthesis, while ORF4 code for an *FNR*-type transcription factor, related to an O_2_-responsive regulator of *hmp* expression, able to repress the Hmp-mediated NO detoxification pathway of *E. coli*.

Thus, this discovery, that other major classes of antibiotics may induce the SOS response could have worthwhile implications for antibiotic stewardship efforts in preventing the emergence of resistance.

## Introduction

*Escherichia coli* is a well-known commensal of the gastrointestinal tract of vertebrates, including humans (Tenaillon et al., 2010), but several strains may cause also enteric and extra-enteric diseases such as urinary tract infection or sepsis (Kaper et al., 2004). *E. coli* is in the fluoroquinolone’s spectrum of action, and these antibiotics are widely used to treat such infections (Lode, 2014; Rice, 2012). Historically, fluoroquinolone resistance was found to develop solely through chromosomal-mediated mechanisms, but plasmid-mediated quinolone resistance (PMQR) genes are now being identified more and more frequently in clinical isolates (Strahilevitz et al., 2009). *qnr* genes are important PMQR determinants, with six families described so far (*qnrA*, *qnrB*, *qnrC*, *qnrD*, *qnrS* and *qnrVC*) (Ruiz, 2019).

Among these *qnr* genes, *qnrD* was first described in *Salmonella enterica* isolates located on a 4,270 bp-long non conjugative plasmid (p2007057), a very different context from that of the other *qnr* genes in terms of plasmid size (Cavaco et al., 2009). Soon after, we reported for the first time the presence of smaller *qnrD*-plasmids (∼2.7 kb, with pDIJ09-518a as the archetype) in several bacteria belonging to *Morganellaceae* and proposed, what is now considered as the likely scenario, that the origin of *qnrD* lies within an as-yet-unidentified progenitor from this family (Guillard et al., 2014; Ruiz, 2019). pDIJ09-518a is a 2,683 bp-long plasmid harbouring four open reading frames (ORFs), including *qnrD*, and exhibits only 53% identity with the plasmid found in *S. enterica* (Fig. S1) (Guillard et al., 2012). No function has yet been found for ORF2, ORF3 or ORF4. *qnrD*-plasmids can be roughly divided into two categories: the pDIJ09-518a-like plasmids (∼2.7 kb) and the p2007047-like plasmids (∼4.3 kb). To date, among the 53 fully sequenced *qnrD*-plasmids, ∼81% are reported to be pDIJ09-518a-like plasmids and ∼19% to be p2007057-like plasmids (Supplementary Table S3 and Fig. S1).

The SOS stress response is a key mechanism by which bacteria respond to DNA damage (Baharoglu and Mazel, 2014; Erill et al., 2007). Oxidative stress and sub-inhibitory concentrations (sub-MIC) of antibiotics can cause DNA damage, triggering the SOS response (Baharoglu et al., 2013; Baharoglu and Mazel, 2011; Kohanski et al., 2010). Once triggered, the SOS response favours bacterial survival in numerous ecological settings, including the likely emergence of antibiotic resistant isolates in patients, when sub-MIC antibiotic concentrations occur at the infection site, through the transient increase of mutation rate accompanying the SOS response (Matic, 2019). Fluoroquinolones are known to induce the SOS response in *E. coli* (Baharoglu and Mazel, 2014; Recacha et al., 2017). In contrast, aminoglycosides are able to induce the SOS response in bacteria such as *Vibrio cholerae* but not in *E. coli*, which is better equipped to face oxidative stress (Baharoglu et al., 2014, 2013; Baharoglu and Mazel, 2011). Two main mechanisms have been reported to explain why aminoglycoside-mediated oxidative stress in *E. coli* does not lead to SOS induction, and include: (i) the production of the general stress response sigma factor RpoS, which induces DNA polymerase IV, and (ii) increased activity of the GO-repair system, which removes the mutagenic oxidized guanine (GO lesion) (Michaels and Miller, 1992), along with the base excision repair (BER) pathway (Baharoglu et al., 2013). Nitric oxide (NO), which can be converted to other reactive nitrogen species (RNS), can also cause DNA damage, inducing also the SOS response (Lobysheva et al., 1999; Nakano et al., 2005a). But, *E. coli* possesses the well characterized Hmp flavohaemoprotein that detoxifies NO either by producing nitrate (NO_3_^-^) under aerobic conditions, or by a O_2_ autoreduction to nitrous oxide (N_2_O) (Cruz-Ramos et al., 2002; Stevanin et al., 2007).

In this study, we show that in *E. coli* carrying pDIJ09-518a-like plasmids the SOS response can nonetheless be triggered by sub-MIC levels of aminoglycosides through NO accumulation. NO accumulation is all at once due to higher NO formation and to the repression of the Hmp-mediated detoxification pathway driven by proteins encoded onto this small *qnrD*-plasmid.

## Results

### *qnrD* expression is SOS regulated and triggered by aminoglycosides

Transcription of the *qnrB and qnrD* genes has been shown to be controlled by the LexA-mediated SOS response. Unlike *qnrB*, the regulation of the most recently characterized PMQR gene, *qnrD,* by the SOS response has only been partially described, given that LexA dependence has not yet been evidenced (Briales et al., 2012; Re et al., 2009). In *E. coli*, nucleofilament of the SOS activator RecA induce the LexA repressor self-cleavage, therefore inducing the SOS global regulatory network (Baharoglu and Mazel, 2014). All the genes belonging to the SOS regulon carry a SOS-box in their promoter. The SOS-box, alternatively named LexA-box, is a 16 bp-long sequence recognized by the LexA repressor. To fill the knowledge gaps regarding *qnrD* regulation, we conducted a multiple alignment analysis of all the 53 *qnrD*-plasmids, fully sequenced and available in GenBank (Supplementary Table S3).

Among these plasmids, we found a highly conserved putative SOS-box upstream of the *qnrD* start codon (see the highlighted logo consensus sequence in the Fig. 1A). This sequence is similar (13/16 identical bases) to the one found in *E. coli* (Fig. 1A).

**Fig. 1.**
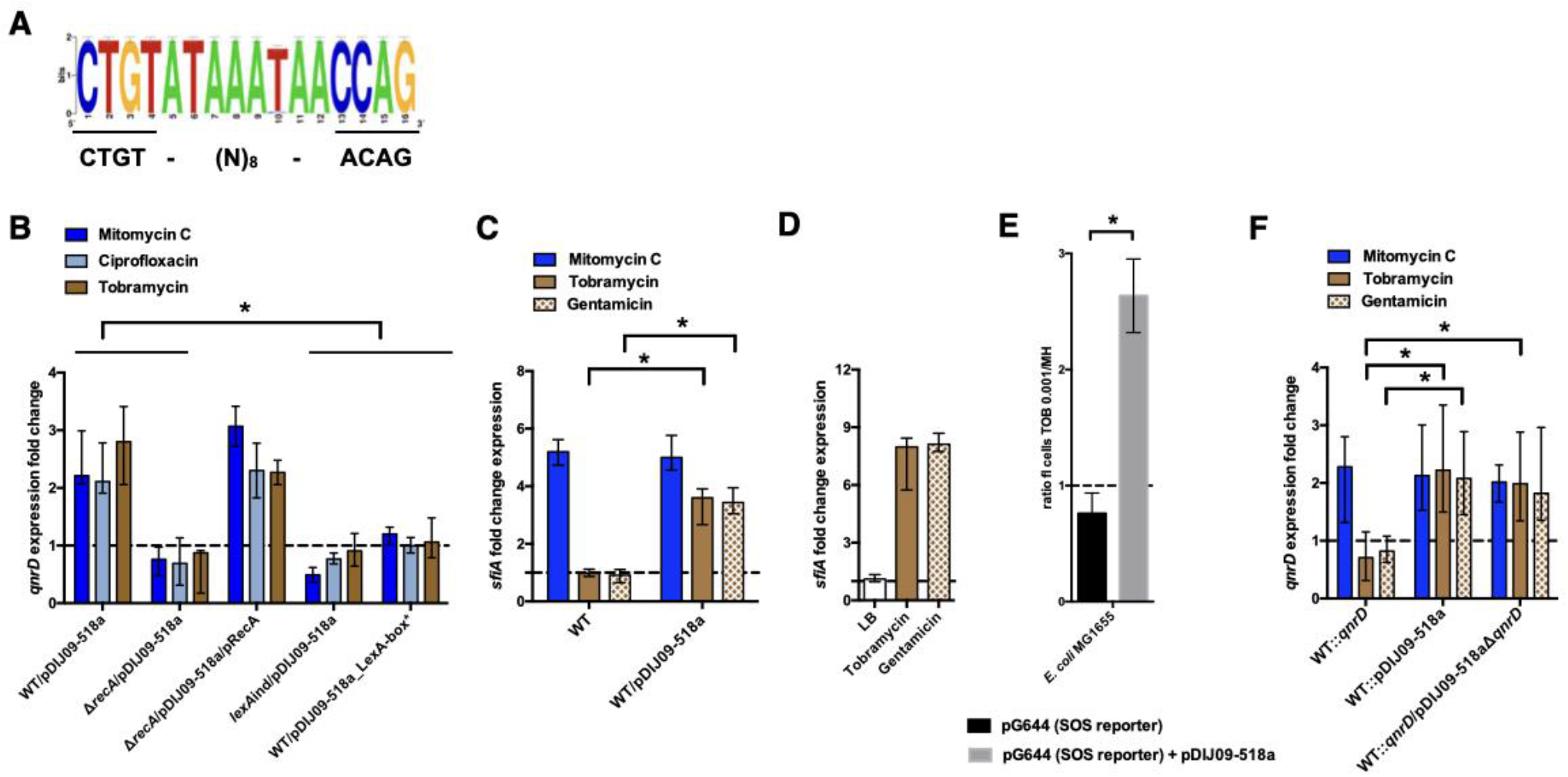
*qnrD* regulation is SOS-mediated and aminoglycosides induce the SOS in *E. coli* because of the *qnrD*-plasmid backbone. **(A)** The *qnrD* SOS-box conservation by visualization of the consensus sequence logo generated from the 53 fully *qnrD*-plasmid sequences. The consensus sequence for *E. coli* is indicated below. **(B)** Relative expression of *qnrD* in *E. coli* MG1656 (WT) derived isogenic strains carrying pDIJ09-518a or pDIJ09-518a with a modified *qnrD*-SOS-box (LexA-box*), exposed to mitomycin C (dark blue), ciprofloxacin (blue) or tobramycin (brown) in comparison to expression in LB, normalized with *dxs*. **(C)** Relative expression of *sfiA* in *E. coli* MG1656 (WT) with or without pDIJ09-518a, exposed to mitomycin C (dark blue), tobramycin (brown) or gentamicin (dotted brown) in comparison to expression in LB, normalized with *dxs*. **(D)** Relative expression of *sfiA* in *E. coli* MG1656 (WT) with pDIJ09-518a, grown in LB, or either with tobramycin or with gentamicin, in comparison to the expression in *E. coli* MG1656, in the three culture conditions, and normalized with *dxs*. (E) Histogram bars show the ratio of GFP fluorescence in a *E. coli* MG1655 WT carrying or not the pDIJ09-518a plasmid in the presence of tobramycin (0.001 μg/mL) over fluorescence of the same strain grown in LB reflecting induction of SOS. Black bars stand for strain with the SOS reporter vector and grey bars stand for strain carrying the *qnrD*-plasmid pDIJ09-518a and the SOS reporter vector. **(F)** Relative expression of *qnrD* in *E. coli* MG1656 with *qnrD* and its own promoter, or the native *qnrD*-plasmid inserted into the chromosome, or chromosomal *qnrD* complemented with pDIJ09-518aΔ*qnrD*, exposed to mitomycin C (dark blue), tobramycin (brown) or gentamicin (dotted brown) in comparison to expression in LB, normalized with *dxs*. Data represent median values of 6 independent biological replicates (2 replicates Δ*recA*/pDIJ09-518a/pRecA), and error bars indicate upper/lower values. \**P* <0.05 Wilcoxon matched-pairs signed-rank test.

We demonstrated the functionality of this SOS-box and its dependence on the RecA and LexA proteins, which are both essential for the SOS response. To do this, we quantified *qnrD* gene expression levels from the native *qnrD*-plasmid, pDIJ09-518a, in the presence of sub-MIC levels of mitomycin C and ciprofloxacin, two well-known SOS-inducers (Supplementary Table 2). It has been clearly established that aminoglycosides do not induce the SOS response in *E. coli* (Baharoglu et al., 2014, 2013; Baharoglu and Mazel, 2011) and therefore the aminoglycoside tobramycin was used as a negative control in our qRT-PCR assays. Using *E. coli* MG1656/pDIJ09-518a grown in LB (MG1656 henceforth referred to as *E. coli* in the text and WT in the figures), we found a 2.18 and 2.02-fold increase in *qnrD* expression induced by mitomycin C and ciprofloxacin, respectively. Unexpectedly, upon tobramycin treatment, we found a 2.8-fold increase in *qnrD* expression (Fig. 1B). To confirm the role of the SOS in induced *qnrD* transcription in the presence of these three drugs, *qnrD* RNA levels were assessed in isogenic *E. coli/*pDIJ09-518a derivatives where the SOS response was not effective due to (i) deletion of its activator (Δ*recA*), (ii) a mutation leading to a non-cleavable repressor (*lexAind*) or (iii) inactivation of the *qnrD* SOS-box directly on pDIJ09-518a (LexA-box*: wild type sequence: <u>CTG</u>TATAAATAACCAG; modified SOS-box: <u>AGC</u>TATAAATAACCAG) (Fig. 1B). As expected, in strains where the SOS response was blocked, *qnrD* expression was not increased by ciprofloxacin or mitomycin C treatments, but this response was also blocked in the presence of tobramycin. To conclusively assert that RecA is needed to increase *qnrD* expression by SOS-dependent regulation, we showed that *qnrD* expression upon tobramycin exposure was increased in the *recA* mutant complemented strain, comparable to the levels detected in the *E. coli*/pDIJ09-518a strain (Fig. 1B).

As shown in figure S2, by determining the growth of both *E. coli* and *E. coli* carrying the *qnrD*-plasmid, we showed that sub-MIC tobramycin treatment has no impact on the viability of *E. coli*/pDIJ09-518a.

To further explore our finding that the SOS response could be triggered by aminoglycosides in *E. coli* carrying pDIJ09-518a, we quantified the expression of *sfiA,* a well-known SOS regulon-induced gene (Huisman and D’Ari, 1981), in the presence of aminoglycosides. As shown in figure 1C, neither tobramycin nor gentamicin were able to increase *sfiA* expression in *E. coli*. In *E. coli* harbouring pDIJ09-518a, however, a 3.13- and 3.71-fold change in *sfiA* expression was measured in the presence of sub-MICs of tobramycin and gentamicin, respectively, confirming that aminoglycosides can induce the SOS response in *qnrD*-plasmid-bearing *E. coli* strains. We also confirmed that pDIJ09-518a did not induce the SOS-response in the absence of aminoglycosides in *E. coli* (Fig. 1D).

To corroborate SOS-mediated *qnrD* expression with exposure to aminoglycosides, we looked at SOS induction in *E. coli* (MG1655 for these experiments) using a previously published SOS reporter setup (Baharoglu et al., 2010). In this system, a GFP-encoding gene is put under the control of the well characterized SOS-driven *recN* promoter (plasmid pG644, Table S3) (Baharoglu et al., 2010), and GFP fluorescence, measured by flow cytometry, gives a readout of the SOS induction level. Using this, we confirmed that, in the presence of aminoglycosides, the SOS response was induced in *E. coli* MG1655 carrying pDIJ09-518a (2.6-fold higher), while no increase in fluorescence was observed in WT *E. coli* MG1655 (Fig. 1E).

### The plasmid backbone contributes to the increase of the *qnrD* gene expression upon aminoglycosides exposure

To determine which components of the pDIJ09-518a plasmid contributed to the SOS response induction in the presence of sub-MIC aminoglycoside treatments, we inserted into the chromosome of *E. coli*, in the *cynX* and *lacA* chromosome intergenic region, either the *qnrD* gene with its own promoter (WT::*qnrD*) alone or the entire plasmid (WT::pDIJ09-518a). For both strains, *qnrD* expression was increased (2.27- and 2.13-fold) in response to treatment with mitomycin C, confirming that neither *qnrD* nor pDIJ09-518a insertion into the *E. coli* chromosome had a negative effect on the SOS induction pathway (Fig. 1F). However, in the presence of sub-MIC levels of tobramycin and gentamicin, no change in *qnrD* transcripts was found in *E. coli* WT::*qnrD,* whereas when the entire plasmid was inserted into the chromosome, *qnrD* transcript levels were increased upon both tobramycin and gentamicin exposure (2.22- and 2.08-fold increase, respectively) (Fig. 1F). Finally, in a WT::*qnrD* strain complemented with the *qnrD*-deleted-pDIJ09-518a plasmid (pDIJ09-518aΔ*qnrD*), exposure to sub-MIC levels of tobramycin or gentamicin induced *qnrD* transcription from the chromosomal site to levels similar to those observed for the WT::pDIJ09-518a strain. Altogether, these results confirmed that in a *qnrD*-harbouring *E. coli* strain, the pDIJ09-518a plasmid backbone without the *qnrD* gene is sufficient to elicit an SOS response by aminoglycosides.

### Small *qnrD*-plasmid promotes nitrosative stress in *E. coli*

In *E. coli*, reactive oxygen species (ROS) are well-known inducers of the SOS response, but as mentioned above, exposure to sub-MIC of aminoglycosides does not lead to ROS accumulation because of the high-level stability of the RpoS (Baharoglu et al., 2013). We hypothesized that carrying *qnrD*-plasmid pDIJ09-518a could affect this stability and therefore increase ROS, causing SOS induction. Thus, we tested the effect of tobramycin on oxidative stress in *E. coli* or *E. coli*/pDIJ09-518a (Fig. S3). Dihydrorhodamine 123 (DHR) oxidation detects the presence of hydrogen peroxide (H_2_O_2_), resulting in the dismutation of the superoxide anion, which is reduced into hydroxyl radicals according to the Fenton reaction (Henderson and Chappell, 1993). Then, fluorescence is measured as an indicator of ROS generation. For this assay, ciprofloxacin was used as a positive inducer of ROS formation. We found no ROS formation induced by tobramycin in either *E. coli* or *E. coli*/pDIJ09-518a (Fig. 2A, ratio ∼ 1, brown bar). As previously reported by Machuca *et al*. for other PMQR determinants (Machuca et al., 2014), we did not find increased ROS formation in *E. coli* carrying the *qnrD*-plasmid upon exposure to sub-MIC of ciprofloxacin (Fig.2A, ratio ∼ 1, blue bar). However, the underlying mechanism explaining these findings has yet to be identified.

**Fig. 2.**
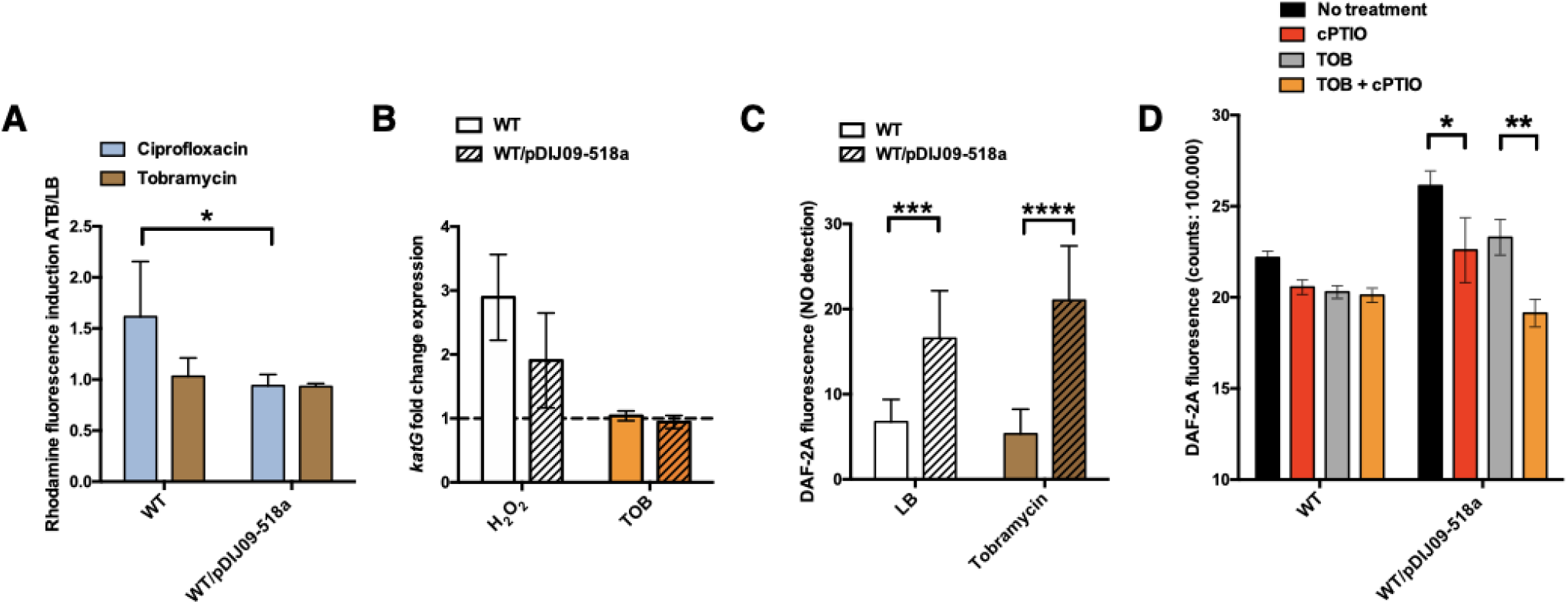
Small *qnrD*-plasmid promotes nitrosative stress in *E. coli*. **(A)** ROS formation for *E. coli* MG1656 (WT) and its derivative carrying pDIJ09-518a cultured in LB or exposed to tobramycin. Production of ROS was calculated as the mean ratio of the DHR-123 fluorescence of the treated samples to the control samples (3 independent biological replicates for each condition). Data were analysed using a 2-way ANOVA with an *P* value < 0.05 for strains as a source of variation in the overall ANOVA. \**P* <0,05 using Tukey’s multiple comparisons test. Mean difference for WT compared to WT/pDIJ09-518a exposed to ciprofloxacin was 0.6753; 95% CI of difference [0.07565; 1.294]. Error bars represent the SD. **(B)** Relative expression of *katG* in *E. coli* MG1656 (WT) and in *E. coli* MG1656 carrying pDJJ09-518a, in comparison to expression in LB, normalized with *dxs*. Data represent median values of 2 independent biological replicates and error bars indicate upper/lower values. Wilcoxon matched-pairs signed-rank test. **(C)** NO formation for *E. coli* MG1656 (WT) and its derivative carrying pDIJ09-518a culture in LB or exposed to tobramycin. Production of NO was calculated as the mean ratio of the DAF-2A fluorescence (2 independent biological replicates for each condition, n = 9 measures). Data were analysed using a 2-way ANOVA with a P value < 0.0001 for strains as a source of variation in the overall ANOVA. \*\*\**P* < 0.001 and \*\*\*\**P* < 0.0001 using a Tukey’s multiple comparisons test. The mean difference for WT compared to WT/pDIJ09-518a grown in LB was −9.790 [95% CI, −15.75; −3.825]. The mean difference for WT compared to WT/pDIJ09-518a exposed to tobramycin was −15,66 [95% CI, −21.62; −9.695]. Error bars represent the SD. Data represent median values of 4 independent biological replicates, and error bars indicate upper/lower values. Wilcoxon matched-pairs signed-rank test. **(D)** Histogram bars show the DAF-2A fluorescence, in *E. coli* WT and WT/pDIJ09-518a as a measure of intracellular in NO obtained using a FACS-based approach, with or without NO scavenger (Carboxy-PTIO, cPTIO). Data were analysed using a 2-way ANOVA with a P value < 0.0001 for treatment as a source of variation in the overall ANOVA. \**P* < 0.05 and \*\**P* < 0.01 using a Tukey’s multiple comparisons test. The mean difference for WT/pDIJ09-518a grown in LB compared to WT/pDIJ09-518a grown in LB and cPTIO was 3.545 [95% CI, 0.4866; 6.603]. The mean difference for WT/pDIJ09-518a grown with Tobramycin (TOB) compared to WT/pDIJ09-518a grown with Tobramycin (TOB) and cPTIO was 4.160 [95% CI, 1.163; 7.157]. Error bars represent the SD. Data represent median values of 5 independent biological replicates, and error bars indicate upper/lower values.

To confirm results from this ROS-detecting dye-based assay, we quantified *katG* expression as an H_2_O_2_ transcriptional reporter. We found increased expression of *katG* induced by H_2_O_2_ in both the WT strain or plasmid-bearing *E. coli*/pDIJ09-518a (Fig. 2B, ratio ∼ 3 and 2, white bars). But interestingly, we did not find any increased *katG* expression induced by tobramycin in either *E. coli* or *E. coli*/pDIJ09-518a (Fig. 2B, ratio ∼ 1, orange bars), confirming that SOS induction in *E. coli* carrying the *qnrD*-plasmid exposed to tobramycin is not due to increased intracellular ROS.

In an effort to find another SOS inducer, we measured the intracellular production of nitric oxide (NO) by determining 5,6-Diaminofluorescein diacetate (DAF-2 DA) fluorescence (Fig. S3) (Kojima et al., 1998; Lobysheva et al., 1999). Strikingly, NO production was significantly elevated in the strains carrying the *qnrD*-plasmid in the absence of an aminoglycoside stimulus, and only increased slightly further when the antibiotic was added at sub-MIC levels (16.55 when strains were grown in LB and an increase of up to 21.01 in the presence of tobramycin) (Fig. 2C). To confirm that the NO production quantified in *E. coli*/pDIJ09-518a was due to the plasmid carriage, we quantified NO production with the addition of a NO scavenger (Carboxy-PTIO, cPTIO) (Fig. 2D). NO levels were decreased in the presence of cPTIO. These results strongly suggest that the carriage of the *qnrD* plasmid by itself induces a nitrosative stress in *E. coli*.

### The GO-repair system is involved in the aminoglycoside-induced SOS response in *E. coli* carrying small *qnrD*-plasmid

Considering that the *qnrD*-plasmid-mediated NO and the aminoglycosides seem to be prerequisites to the SOS response induction in *E. coli*/pDIJ09-518a, we considered deleterious effects common to both these stimuli that could induce the SOS response. Aminoglycosides, as well as NO and other reactive nitrogen species, such as dinitrogen trioxide (N_2_O_3_) and peroxynitrite (ONOO^-^), form 8-oxo-G (7, 8-dihydro-8-oxoguanine) (Baharoglu et al., 2013; Nakano et al., 2005b, 2005a). It has been described that the incorporation of 8-oxo-G into DNA leads to A/G mismatch during replication (Grollman and Moriya, 1993; Michaels et al., 1992; Michaels and Miller, 1992). To prevent or repair these types of oxidative lesions, *E. coli* uses the GO-repair system, with MutT and MutY as two of the key players in this error-prevention system (Baharoglu et al., 2014; Boiteux et al., 2017; David et al., 2007; Foti et al., 2012; Grollman and Moriya, 1993; Maki and Sekiguchi, 1992; Michaels et al., 1992). Incomplete action of the GO-repair system leads to the formation of double strand DNA breaks (DSBs), which are lethal if unrepaired (Foti et al., 2012). MutT acts as a nucleotide pool sanitizer removing 8-oxo-G triphosphate, while MutY removes the adenine base from 8-oxo-G/A mispairing. We further hypothesized that the SOS response could rely on the deleterious effect of unremoved oxidized guanines due to the combined stress of NO and aminoglycosides.

To test the hypothesis that the SOS response induced by aminoglycosides is linked to 8-oxo-G-mediated DSBs formation because of their incomplete removal, we measured the SOS response in *E. coli* carrying pDIJ09-518a with or without overexpression of MutT. As shown in Fig. 3A we found that the SOS response, measured through *sfiA* transcription levels, was induced in the presence of aminoglycosides in *E. coli*/pDIJ09-519a carrying the empty vector used for MutT overexpression (3.15-fold increase), while it was not increased in *E. coli*/pDIJ09-518a over-expressing MutT (0.59-fold change).

**Fig. 3.**
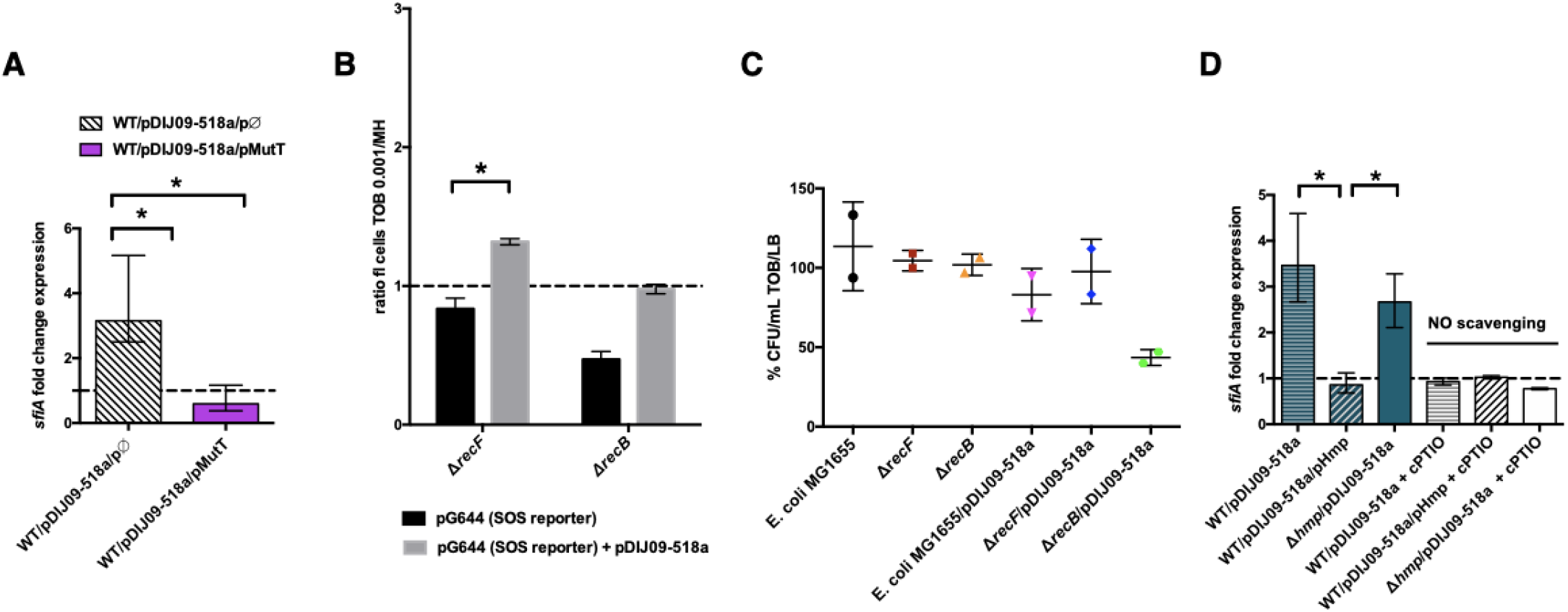
Aminoglycosides induce SOS in *E. coli*/pDIJ09-518a due to overwhelmed GO-repair pathway associated with inactivated Hmp. **(A and D)** Relative expression of *sfiA* in *E. coli* MG1656 (WT) isogenic strains carrying pDJJ09-518a, overexpressing the GO-repair system protein MutT and the *hmp*-deleted mutant, exposed to tobramycin, treated with the NO scavenger carboxy-PTIO (cPTIO) (for D), in comparison to expression in LB, normalized with *dxs*. Data represent median values of 6 independent biological replicates (2 replicates for cPTIO condition) and error bars indicate upper/lower values. Wilcoxon matched-pairs signed-rank test. **(B)** Histogram bars show the ratio of GFP fluorescence in a *E. coli* MG1655 *ΔrecB* and *ΔrecF* in the presence of tobramycin (0.001 μg/mL) over fluorescence of the same strain grown in LB reflecting induction of SOS. Black bars stand for strain with the SOS reporter vector and grey bars stand for strain carrying the *qnrD*-plasmid pDIJ09-518a and the SOS reporter vector. **(C)** Impact of *recB* gene inactivation in *E. coli* harbouring the *qnrD* plasmid on growth in sub-MIC tobramycin. Histogram bars represent the percentage of the ratio of CFU/ml for each strain in tobramycin (0.001 µg/mL) over CFU/mL in LB. Data represent median values of 2 independent biological replicates, and error bars indicate standard deviation. Wilcoxon matched-pairs signed-rank test.

To further determine if the incomplete action of the GO-repair system may lead to the accumulation of DSBs in *E. coli*/pDIJ09-518a exposed to aminoglycosides, we used two *E. coli* MG1655 strains wherein the SOS could either be induced or not activated in the presence of DSBs (*E. coli* Δ*recF* and *E. coli* Δ*recB*, respectively). The RecFOR pathway allows RecA nucleo-filament formation on ssDNA breaks, whereas the RecBCD recruits RecA onto DSBs (Baharoglu and Mazel, 2014; Kuzminov, 1999). As shown in figure 3B, SOS was not induced by aminoglycosides in the *recB* deleted-strain, whereas a slight induction of the SOS response was detected in the *recF* deletant (1.4-fold change compared to growth in LB). These results suggest that in *E. coli* carrying *qnrD*-plasmids, DSBs are produced upon aminoglycoside treatment inducing the SOS response necessary for bacterial survival. The deletion of *recB* in *E. coli* harbouring the plasmid causes loss of viability after exposure to tobramycin as compared to the wild-type *E. coli* MG1655 or the *recF* mutant (Fig. 3C).

### Role of the NO-detoxifying Hmp in the SOS induction upon exposure to sub-MIC of aminoglycosides

In *E. coli*, the Hmp flavohaemoprotein has been described as the key player in detoxifying NO under aerobic conditions (Poole et al., 1996). We therefore hypothesized that NO accumulation could be due to ineffective Hmp-mediated detoxification leading to the accumulation of 8-oxo-G and 8-nitro-G DNA lesions and thereby, an SOS response induction in *E. coli* exposed to aminoglycosides. To test this hypothesis, we over-expressed Hmp in *E. coli*/pDIJ09-518a. As shown in Fig. 3D, the *sfiA* transcription increase observed in *E. coli*/pDIJ09-518a in the presence of aminoglycosides (3.14-fold increase over growth without aminoglycosides) was now abolished when Hmp was over-expressed (0.85-fold decrease compared to growth without aminoglycosides). To confirm this finding, we quantified the SOS induction in a *qnrD*-plasmid-harbouring *E. coli* strain where *hmp* had been deleted. In this strain, unable to detoxify NO, the carriage of the *qnrD*-plasmid increased the expression of *sfiA* (∼ 2.5-fold) when exposed to tobramycin (Fig. 3D). We confirmed that neither deleting *hmp* nor carrying the empty vector used to over-express Hmp, increased the expression of *sfiA* in *E. coli* grown in LB medium without aminoglycosides (Fig. S4A). Furthermore, the addition of tobramycin did not induce the SOS response in the *hmp* mutant (Fig. S4B). After exposure to tobramycin, complementation of the *hmp* mutant carrying the *qnrD*-plasmid with pHmp (a plasmid expressing *hmp*), did not trigger the SOS response, whereas the SOS was induced in the same genetic context but no *hmp* complementation (Fig. S4B). To confirm our hypothesis, we quantified SOS induction in the same derivative strains exposed to aminoglycosides using a NO scavenger (cPTIO) assay. It is noteworthy that in the case without any NO, the SOS response was not induced (Fig. 3D).

Our results show that *qnrD-*plasmid-bearing *E. coli* undergoes nitrosative stress because of a much less effective Hmp-mediated detoxification that leads to SOS response induction when exposed to aminoglycosides. These findings underscore that both NO accumulation and tobramycin are needed for the induction of the SOS response in *E. coli* carrying pDIJ09-518a.

### Small *qnrD*-plasmid genes promote NO formation and inhibit NO detoxification

Next, we tried to establish which pDIJ09-518a ORF(s) promoted the NO accumulation leading to the tobramycin-induced SOS response in *E. coli*. Among bacteria, NO synthesis by NO synthase has been seldomly reported, *i.e.* in *Nocardia spp*, though this bacterial enzyme is different from the mammalian version. In most other bacteria, however, the main sources of NO come from the activity of nitrite and nitrate reductases, which catalyse the reduction of nitrate (NO_3_^-^) and/or nitrite (NO_2_^-^) to NO (Crane et al., 2010). Querying the pBLAST/psiBLAST databases, we found that ORF3 encoded a flavin adenine dinucleotide (*FAD*)-binding oxidoreductase with a NAD(P) binding Rossmann-fold (36% protein identity over 43% of the FAD-binding oxidoreductase domain from an *Erythrobacter* spp protein) that could be involved in NO production. We further found that the adjacent ORF4 encoded a cAMP-receptor *protein* (*CRP*)/fumarate and nitrate reduction regulatory protein (*FNR*)-type transcription factor (CRP/FNR; 34% protein identity over 71% of CRP/FNR from *Pedobacter panaciterrae*), an O_2_-responsive regulator of *hmp* expression.

We hypothesized that ORF3 could be involved in the aminoglycoside-induced SOS response by promoting NO production, which was tested by the deletion of ORF3 from pDIJ09-518a (*E. coli*/pDIJ09-518aΔORF3). ORF3 deletion led to the loss of *sfiA* induction in response to tobramycin (Fig. 4A, brown bar). Complementation by ORF3 (*E. coli*/pDIJ09-518aΔORF3/pORF3) restored the SOS response (Fig. 4A). In addition, we showed that the deletion of ORF3 decreased NO production compared to that measured in the presence of the native *qnrD*-plasmid in *E. coli* (Fig. 4B).

**Fig. 4.**
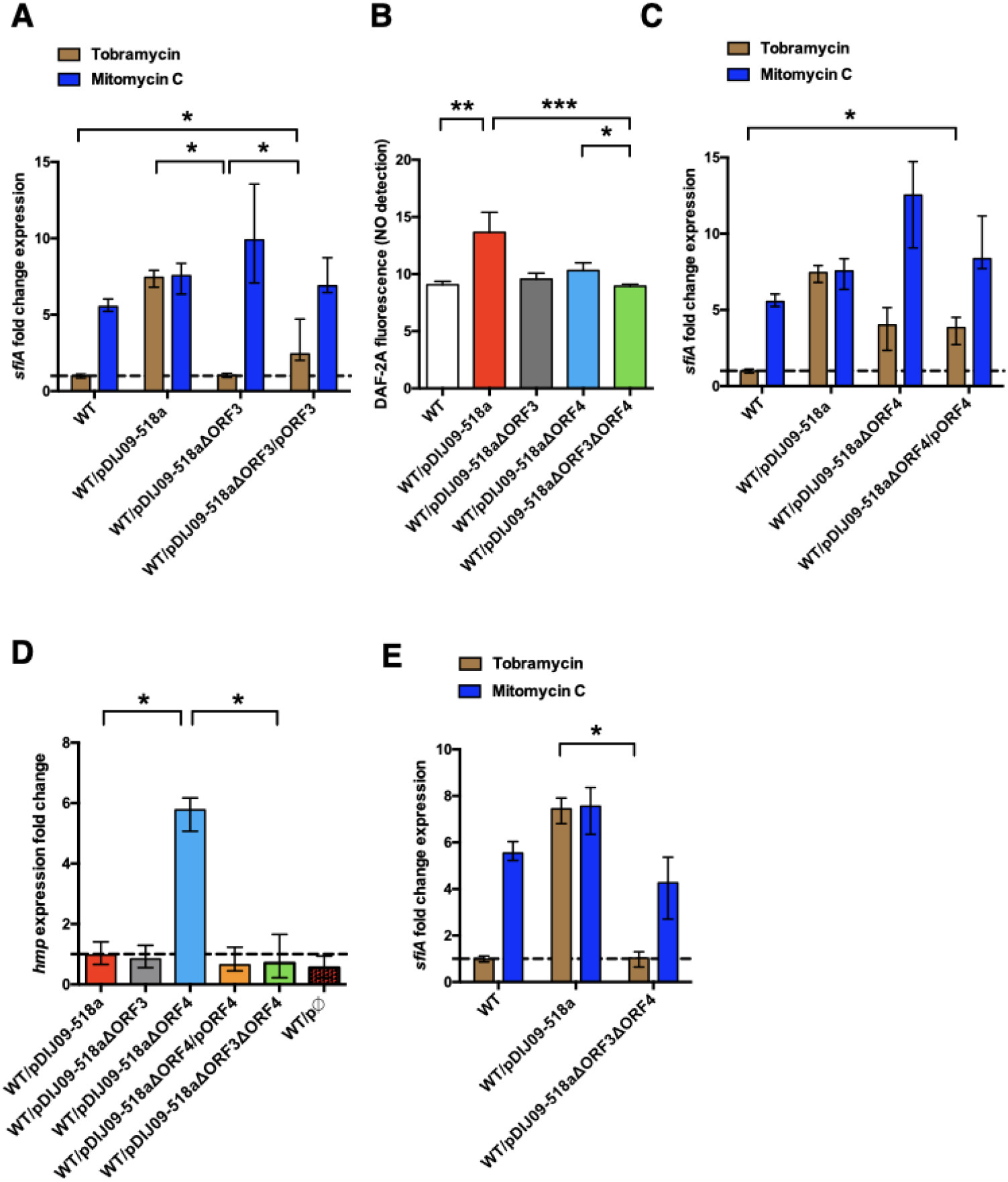
Deletion of ORF3 decreases the SOS response induction, after tobramycin treatment and ORF4 regulates the Hmp nitric oxide detoxification pathway. **(A, C and E)** Relative expression of *sfiA* in *E. coli* MG1656 (WT) derived isogenic strains carrying pDIJ09-518a with ORF3 and/or ORF4 deleted and complemented, exposed to mitomycin C (dark blue) or tobramycin (brown) in comparison to expression in LB, normalized with *dxs*. Data represent median values of 6 independent biological replicates, and error bars indicate upper/lower values. \**P* <0.05 Wilcoxon matched-pairs signed-rank test. **(B)** NO formation for the isogenic strains (n=9). Data were analysed using a Kruskal-Wallis test, with a *P* value < 0.0001 for the overall ANOVA. NO formation for each strain was analysed using Dunn’s multiple comparisons test. \**P* < 0.05, \*\**P* < 0.01 and \*\*\**P* < 0.001. Bars represent mean values and SD. **(D)** Relative expression of *hmp* in *E. coli* MG1656 (WT) derivative isogenic strains carrying pDIJ09-518a with ORF3 and/or ORF4 deleted and complemented, or the empty vector, grown in LB, in comparison to expression in *E. coli* MG1656 (WT), normalized with *dxs*. Data represent median values of 6 independent biological replicates, and error bars indicate upper/lower values. Wilcoxon matched-pairs signed-rank test.

To determine the possible effect of the FNR-like O_2_-response regulator encoded by ORF4 in pDIJ09-518a, on *hmp* expression (Cruz-Ramos et al., 2002; Poole et al., 1996), we deleted and complemented this ORF in pDIJ09-518a. We found that ORF4 did not alleviate the SOS response after exposure to aminoglycoside sub-MICs, as measured by *sfiA* expression (Fig. 4C). However, we did find that ORF4 deletion decreased intracellular NO production (Fig. 4B). We next found that ORF4 deletion increased the transcription of *hmp* (Fig. 4D) while *hmp* transcription in *E. coli*/pDIJ09-518aΔORF4 complemented with ORF4 (*E. coli*/pDIJ09-518aΔORF4/pORF4) was similar to that in *E. coli*/pDIJ09-518a. We confirmed that the carriage of the expression vector alone did not impact the transcription of *hmp* compared to the parental *E. coli* in LB (Fig. 4D).

Finally, in the presence of a double deletion in pDIJ09-518a (ΔORF3ΔORF4), we found no SOS induction by tobramycin (Fig. 4E), a slight decrease in NO formation (Fig. 4B) and no *hmp* expression (Fig. 4D), confirming that only ORF4 impacts the level of *hmp* expression. Notably, neither the deletion of ORF3 nor ORF4 alleviated the SOS response induced by mitomycin C (Fig. 4A and 4C, dark blue bars). Significant experimental effort, beyond the scope of this initial report, will be required to biochemically and genetically characterized the activities of purified proteins encoded by the ORF3 and ORF4 genes and their possible role in NO formation and detoxification under aerobic conditions.

## Discussion

It is known that, conversely to *V. cholerae*, sub-MIC of aminoglycosides do not induce the SOS response in *E. coli*. In this study, however, we found and characterize for the first time that the SOS response is induced in *E. coli* carrying a *qnrD*-plasmid upon exposure to aminoglycosides at very low concentration (1% of MIC). This unexpected aminoglycoside-induced SOS response turns to be subsequent to NO accumulation in combination with aminoglycosides that eventually increase 8-oxo-G incorporation into DNA. Thereby, DNA damage appears through DSBs leading to induction to the SOS.

We found that two ORFs of the *qnrD*-plasmid, ORF3 and ORF4, were responsible for this SOS response induction by aminoglycosides in *E. coli*. Indeed, ORF3, which encodes a FAD-binding oxidoreductase, leads to NO production while ORF4, which encodes a CRP/FNR-like protein, inhibits *hmp* expression and thereby hampers the detoxification of NO. Then, we propose a model (Fig.4) of the pathway by which aminoglycosides induce the SOS response in *E. coli* carrying the *qnrD*-plasmid. In the absence of aminoglycosides, *E. coli* can repair the DNA damage resulting from the carriage of the *qnrD* plasmid, and the SOS response is not induced (Fig. 5A). In this case, the GO-repair-system sanitizes efficiently the 8-oxo-G produced by the *qnrD*-plasmid-mediated NO accumulation. However, in the presence of aminoglycoside sub-MICs, (Fig. 5B), the level of oxidized guanine increases. The resultant DNA damage yields genotoxic concentrations of alternate nucleotides that the GO-repair system can no longer sanitize efficiently, leading to induction of the SOS response.

**Fig. 5.**
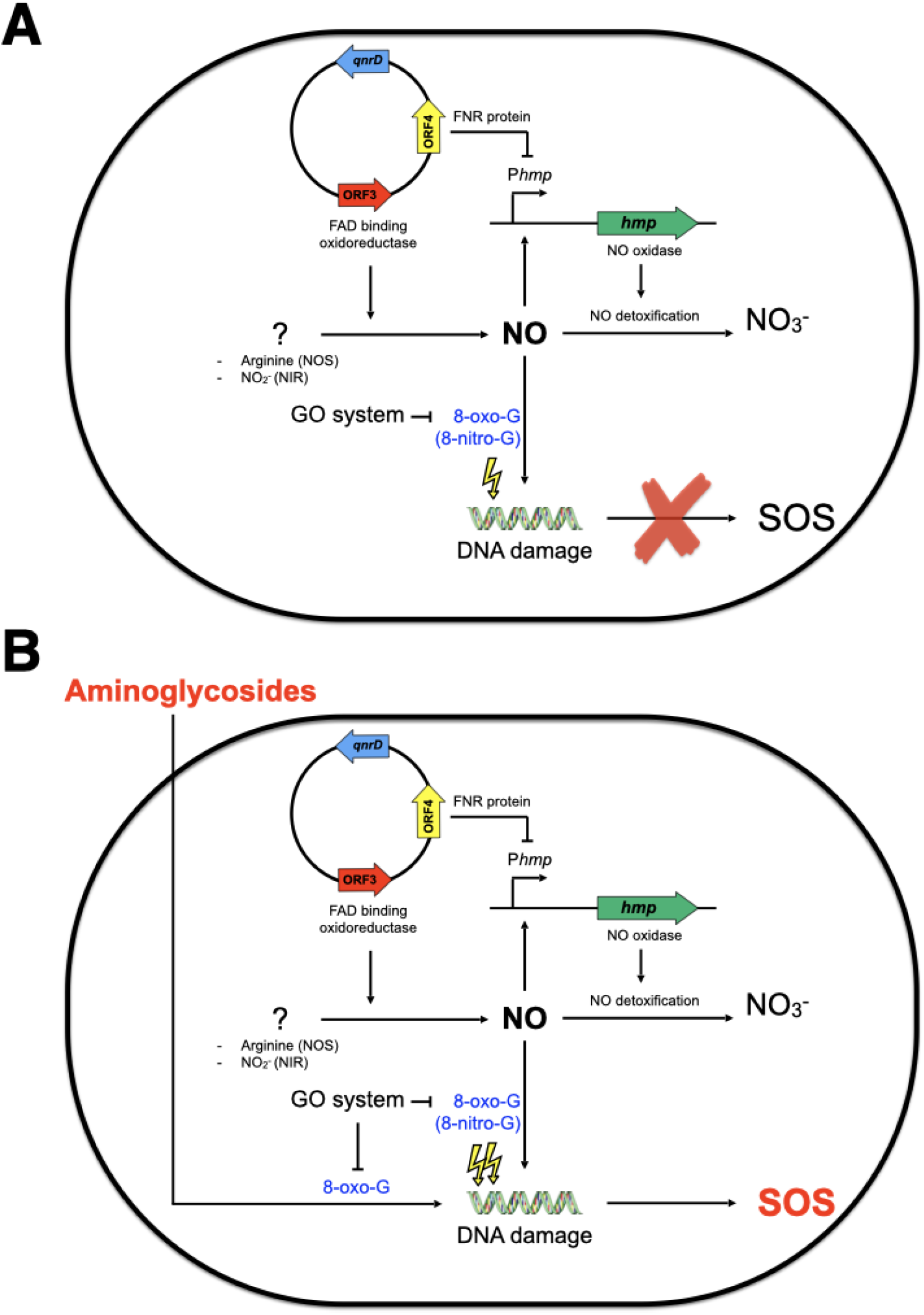
Model of SOS response induction by aminoglycosides in *E. coli* bearing the small *qnrD*-plasmid. Schematic representation of the network leading to SOS induction in *E. coli*/pDIJ09-518a when exposed **(B)** or not **(A)** to sub-inhibitory MIC of aminoglycosides.

Our study shows that the widely accepted lack of SOS response induction in *E. coli* by aminoglycosides may not always be true. When strains bearing the *qnrD*-plasmid are exposed to sub-MICs of aminoglycosides, the SOS response does occur. This is a worrying issue, since we observed that the *qnrD*-plasmid is mobilizable and stable without antibiotic selective pressure (data not shown). Selective pressures maintained by the over-use of antibiotics are the main drivers of resistance. In addition, sub-inhibitory levels of antibiotics (sub-MICs) can select for resistant bacteria and this occurs notably with fluoroquinolones (Andersson and Hughes, 2017). In this regard, expression of Qnr proteins and other PMQR normally confers only low levels of resistance to fluoroquinolones, but they were shown to significantly reduce the bactericidal activity of ciprofloxacin (Allou et al., 2009; Guillard et al., 2013) and to further facilitate selection for higher levels of resistance in Qnr-producing enterobacterial isolates exposed to fluoroquinolones, which can culminate in therapeutic failures (Allou et al., 2009; Guillard et al., 2013; Martínez-Martínez et al., 1998; Robicsek et al., 2006). In addition, it has also recently been shown that *qnrB* promotes DNA mutations and thereby fluoroquinolone-resistant mutants by triggering DNA replication stress.(Li et al., 2019)

In this context, the SOS regulation of such PMQR genes have clinical implications, not only in term of infectious disease treatments, but also to prevent the dissemination of resistance genes. It has been clearly demonstrated that the *qnrB*-mediated quinolone resistance is induced upon exposure to sub-MICs of fluoroquinolone (Re et al., 2009). Therefore, describing for the first-time aminoglycosides that induce the SOS response in *E. coli* carrying a low level of fluoroquinolone resistance determinant (*qnrD*-plasmid) could have worthwhile therapeutic implications by increasing the odds of mutations during the SOS induction, since each of these classes of antibiotics are commonly used as first- or second-line treatment.

## Materials and Methods

### Bacterial strains, plasmids, primers and growth conditions

The bacterial strains, plasmid constructs and primers for PCR analysis used in this work are shown in Supplementary Tables 3 and 4 (Blattner et al., 1997; Mount et al., 1972). The *E. coli* Δ*hmp* (KEIO collection) (Baba et al., 2006) was graciously provided by J-M. Ghigo (Pasteur Institute, Paris). This allele was transduced in MG1656, using P1 phage and selected on agar plates with 50 µg kanamycin /mL. Experiments were performed in lysogeny broth (LB) or in minimum medium at 37°C. For genetic selections, antibiotics were added to media at the following concentrations: 100 μg ampicillin /mL, 0.03 and 0.06 μg ciprofloxacin /mL, 50 μg kanamycin /mL, 50 µg streptomycin /mL or 100 amoxicillin µg/mL. For each strain, the MIC of antibiotics was determined twice for each antimicrobial agent using E-test strips (bioMérieux, Marcy l’Etoile, France). The sub-MICs (e.g. 1% of MIC). of specified antibiotics were used to induce the SOS response, as follows (final concentrations [µg/mL]): ciprofloxacin (CIP) 0.06, gentamicin (GM) 0.00125, mitomycin C (MMC) 0.1 and tobramycin (TM) 0.001

### WT::qnrD and WT::pDIJ09-518a strains construction

DNA fragments were generated by PCR in order to amplify the *qnrD* with its own promoter or the pDIJ09-518a from the native plasmid and *cynX* and *lacA* chromosome intergenic region from the MG1656 genomic DNA. The oligonucleotides primers used to amplify the DNA are shown in the Supplementary Table 4. The three PCR fragments were digested with DpnI and purified with a Qiagen PCR Purification Kit (Qiagen). The three PCR products were assembled as one large fragment (5’ *cynX* - insert - *lacA*3’) by Gibson Assembly (New England Biolabs). The assembled DNA was transformed into electrocompetent WT *E. coli* where they recombined into the *E. coli* MG1656 genome. The transformed bacteria were selected using 0.06 µg ciprofloxacin /mL by incubation for 24h, at 37°C. The insertion was verified by colony PCR with four primers pairs: AB09/AB06 (*qnrD*), AB13/AB08 (plasmid), AB06/AB12 (*qnrD*) and AB07/AB16 (plasmid) (Supplementary Table 4). The positive clones were verified by sequencing using AB09/AB12 and AB13/AB16.

### Plasmid constructions

The *recA*, *mutT,* and *hmp* genes with their own promoters were amplified from the *E. coli* MG1656 genome, with the corresponding Forward/Reverse primers shown in Supplementary Table 4. The PCR products were purified and cloned into pCR2.1®, hereafter called pTOPO, (ThermoFisher Scientific, France) to generate pRecA, pMutT, and pHmp, respectively and selected on plates containing 50 µg kanamycin /mL. The same protocol was used for the complementation of the ORF3 and ORF4 genes to generate pORF3 and pORF4. However, these ORFs were amplified from *E. coli*/pDIJ09-518a.

pDIJ09-518aΔ*qnrD* was obtained by PCR amplification of the native pDIJ09-518a plasmid as DNA template excluding the *qnrD* gene, using the primers described in the Supplementary Table 4. The primers were obtained using the NEBuilder Assembly Tool (New England Biolabs). After digestion by DpnI (New England Biolabs) and purification of PCR products (Qiagen), the fragment obtained was transformed into electrocompetent *E. coli* WT (MG1656) or WT::*qnrD*. Transformants were selected on agar plates containing 0.06 µg ciprofloxacin/mL and were analysed by PCR as described above. The same protocol was used to obtain pDIJ09-518aΔORF3, -ΔORF4 and -ΔORF3ΔORF4, using the indicated primers (Supplementary Table 4).

pDIJ09-518a LexA-box* substitution of CGT to AGC, in the LexA-box or SOS-box binding site, was obtained by using a modified Quick-Change II Site-Directed Mutagenesis kit (Agilent). Briefly, we used the primers containing the substitution (Supplementary Table 4), as described by the manufacturer. The elongation temperature used for the amplification was modified to 68°C for 2 minutes. 18 cycles of amplification were sufficient to obtain a proper amount of modified DNA template. After digestion by DpnI and purification of PCR products, the fragment obtained was transformed into *E. coli* competent TOP10 cells (ThermoFisher). Transformants were selected on agar plates containing 0.06 µg ciprofloxacin/mL and analysed by PCR as described above.

### DNA manipulation and genetic techniques

Genomic DNA (gDNA) was extracted and purified using the Qiagen DNeasy purification kit (Qiagen, Courtaboeuf, France). Isolation of plasmid DNA was carried out using the QIAprep Spin Miniprep kit (Qiagen). Gel extractions and purifications of PCR products were performed using the QIAquick Gel Extraction kit (Qiagen) and QIAquick PCR Purification kit (Qiagen). PCR verifying experiments were performed with Go Taq Green Master Mix (Promega, Charbonnières les Bains, France), and PCRs requiring proofreading were performed with the Q5^®^ High-Fidelity DNA Polymerase (New England BioLabs, Evry, France) as described by the manufacturers. Restriction endonucleases DpnI was used per the manufacturer’s specifications (New England BioLabs). All DNA manipulations were checked by DNA sequencing (GENEWIZ, Takeley, England).

### RNA extraction and qRT-PCR

Strains were grown in LB at 37°C with shaking to exponential phase (OD_600_ = 0.5-0.7). Six biological replicates were prepared. One percent of the MIC of indicated antibiotics was then added to the culture for 30 min to allow for induction of the SOS response. One culture was kept as an antibiotic-free control. Five hundred microliters of exponentially growing cells were stabilized in 1 mL of RNAprotect Bacteria Reagent (Qiagen). After treatment of the culture pellet with lysozyme (Qiagen), subsequent RNA extractions were performed using the RNeasy Mini Kit (Qiagen). The gDNA contaminating the samples was removed with TURBO DNA-free Kit (Ambion, ThermoFisher Scientifics) at 37°C, for 30 min. First-strand cDNA synthesis and quantitative real-time PCR were performed with KAPA SYBR® FAST (CliniSciences, Nanterre, France) on the LightCycler 480 (Roche Diagnostics, Meylan, France) using the primers indicated in Supplementary Table 4. Transcript levels of each gene were normalized to *dxs* as the reference gene control. Gene expression levels were determined using the 2^-ΔΔCq^ method (Bustin et al., 2009; Livak and Schmittgen, 2001) in respect to the MIQE guidelines. Relative fold-difference was expressed either by reference to antibiotic free culture or the WT strain in LB. All experiments were performed as six independent replicates (if not, number of replicates are clearly specified in the legends) with all samples tested in triplicate. Cq values of technical replicates were averaged for each biological replicate allowing us to obtain the ΔCq. After exponential transformation of the ΔCq for the studied and the normalized condition, medians and upper/lower values were determined.

### Flow cytometry

Flow cytometry experiments were performed as described (Baharoglu et al., 2010) and repeated at least 3 times on overnight cultures in MH or MH + sub-MIC tobramycin (0.001 µg/mL). Briefly, overnight cultures of *E. coli* MG1655 and *E. coli* MG1655/pDIJ09-518a and their derivatives were prepared in LB broth with or without sub-MIC of tobramycin and diluted 40-fold into PBS (Phosphate-Buffered Saline, Invitrogen). The GFP fluorescence was measured using the Miltenyi MACSQuant device.

### Detection of intracellular ROS by DHR123 and NO by DAF2-DA

Overnight cultures of *E. coli* MG1656 and *E. coli* MG1656/pDIJ09-518a and its derivatives were diluted 100-fold in fresh LB broth. Dihydrorhodamine 123 (DHR123, Sigma) or 5,6 Diaminofluorescein diacetate (DAF-2 DA, Sigma) were added to 5 mL LB to a final concentration of 2.5 x 10^-3^ µM or 10 µM, respectively. Sub-MICs of ciprofloxacin or tobramycin were added for 30 min to the cultures at OD_600_ 0.5-0.7, and when stated, carboxy-PTIO (cPTIO, Enzo Life Sciences, final 5µM). Two hundred microliters of bacterial cultures were then added to 96-well black flat-bottom plates (Corning) in triplicate. The DHR fluorescence was measured at 507 nm/529 nm (excitation/emission wavelength) whereas the DAF-2 DA fluorescence was measured at 491nm/513nm, on a SAFAS Xenius XC (Safas, Monaco). The values used were corrected by subtracting the values from the negative controls. Experiments were repeated for 2 or 3 independent biological replicates at least 3 times.

### Detection of intracellular NO by flow cytrometry

Overnight cultures of *E. coli* MG1655 and *E. coli* MG1655/pDIJ09-518a were diluted 100-fold into fresh LB broth, and growth until an OD_600nm_ of 0.2 was reached. The DAF-2A, the NO scavenger cPTIO (5µM final concentration) and sub-MIC of tobramycin were then added for 30 min to the cultures. Flow cytometry experiments were performed as described (Baharoglu et al., 2010) and repeated at least 5 times. DAF-2A fluorescence was measured using the Miltenyi MACSQuant device.

### Minimum inhibitory concentration (MIC) determination

MICs were determined by E-test (bioMérieux, Marcy l’Etoile, France) in accordance with EUCAST guidelines. Briefly, MICs of nalidixic acid, ciprofloxacin, levofloxacin, and ofloxacin were determined on Mueller-Hinton (MH) agar inoculated with a 0.5 McFarland (∼ 10^8^ CFU per mL) bacterial suspension. MICs were read after incubation for 18 h at 37°C.

### Growth curves

Overnight cultures of *E. coli* and *E. coli*/pDIJ09-518a were diluted 100-fold into LB broth. Growth curves were obtained using an automated turbidimetric system (Bioscreen C, LabSystem) at 37°C with shaking during 24 h. Optical density measurements at 600 nm (OD_600_) were performed every 5 min with 10 s of shaking prior to reading. The OD_600_, corrected with values from the negative controls, and the corresponding Log10 CFU/mL, were used to fit the growth curves of each studied strain. For both strains, biological experiments were performed in triplicate.

### Viability study after tobramycin treatment

Overnight cultures of *E. coli* MG1655 and *E. coli* /pDIJ09-518a and their derivatives strains were diluted 100-fold into LB broth and LB broth supplemented with 0.001 µg/mL tobramycin. Aliquots were plated on LB agar and incubated at 37°C for 24h. The colony forming units (CFU)/mL were counted. The bars represent the percentage of CFU counted after tobramycin treatment over LB. For all strains, biological experiments were performed in duplicate.

### Bioinformatic analysis and statistics

The prevalence and the dissemination of the small *qnrD*-plasmids into *Enterobacterales* were analysed on the National Centre for Biotechnology Information (NCBI) database. The small *qnrD*-plasmids were classified into three groups (*Morganellaceae*, *E. coli* and others *Enterobacterales)* and into two classes, considering the length of known *qnrD*-plasmids (pDIJ09-581a or p2007057-like).

The LexA-box consensus sequence logo was established using WebLogo http://weblogo.berkeley.edu/logo.cgi (Crooks et al., 2004) taking into account the 16nt of SOS-box of all 53 fully sequenced *qnrD*-plasmids.

For qRT-PCR, a Wilcoxon matched-pairs signed rank test was used to compare median of fold changes (Livak and Schmittgen, 2001; Schmittgen and Livak, 2008; Yuan et al., 2006).

For ROS and NO detection, a two-way ANOVA with *ad hoc* tests (Tukey’s multiple comparisons test or a Dunn’s multiple comparisons test) was used to compare the measured values between the different strains and conditions.

For spontaneous mutations rates, a Wilcoxon matched-pairs signed rank test was used to compare median of Rif CFU/ml/ total CFUs counted.

All the tests were performed using GraphPad Prism version 6. Degree of significance are indicated as follows: * *P* < 0.05; ** *P* < 0.01; *** *P* < 0.001; **** *P* < 0.0001.

## Data availability

Source data are provided as a Source Data file. Flow cytometry data have been deposited in FlowRepository as FR-FCM-Z3MR repository ID (http://flowrepository.org/id/RvFrzhOtiB4Hrd9yMMTEF2gAckZvYVa365phD9U0fVTabQb7ib

CDqV8Gzbgb02dm).

## Acknowledgments

We thank Dr Valerian Dormoy for his careful reading of the manuscript and scientific discussion of the results, Dr Christine Terryn of PICT-URCA platform for technical assistance in imaging core facilities, Dr Arnaud Bonnomet from Inserm UMR-S 1250 for their technical assistance and Prof Grace Stockton-Bliard for proofreading the manuscript.

## Materials & Correspondence

To whom correspondence and material requests should be addressed: Thomas Guillard

Laboratoire de Bactériologie–Virologie-Hygiène Hospitalière-Parasitologie-Mycologie, CHU Reims, Hôpital Robert Debré, Avenue du Général Koenig, 51092 Reims Cedex, France., Tel/Fax: +33 3 26 78 32 10 / +33 3 26 78 41 34

## Supplementary information

**Table S1.** Minimum-inhibitory concentrations for antibiotics used as SOS-response inducers.

| Strains name | Tobramycin (µg/L) | Gentamicin (µg/mL) | Ciprofloxacin (µg/mL) |
| --- | --- | --- | --- |
| *E. coli* MG1656 | 0.125 | 0.125 | 0.004 |
| *E. coli* MG1656/pDIJ09-518a | 0.125 | 0.125 | 0.094 |

**Table S2.**
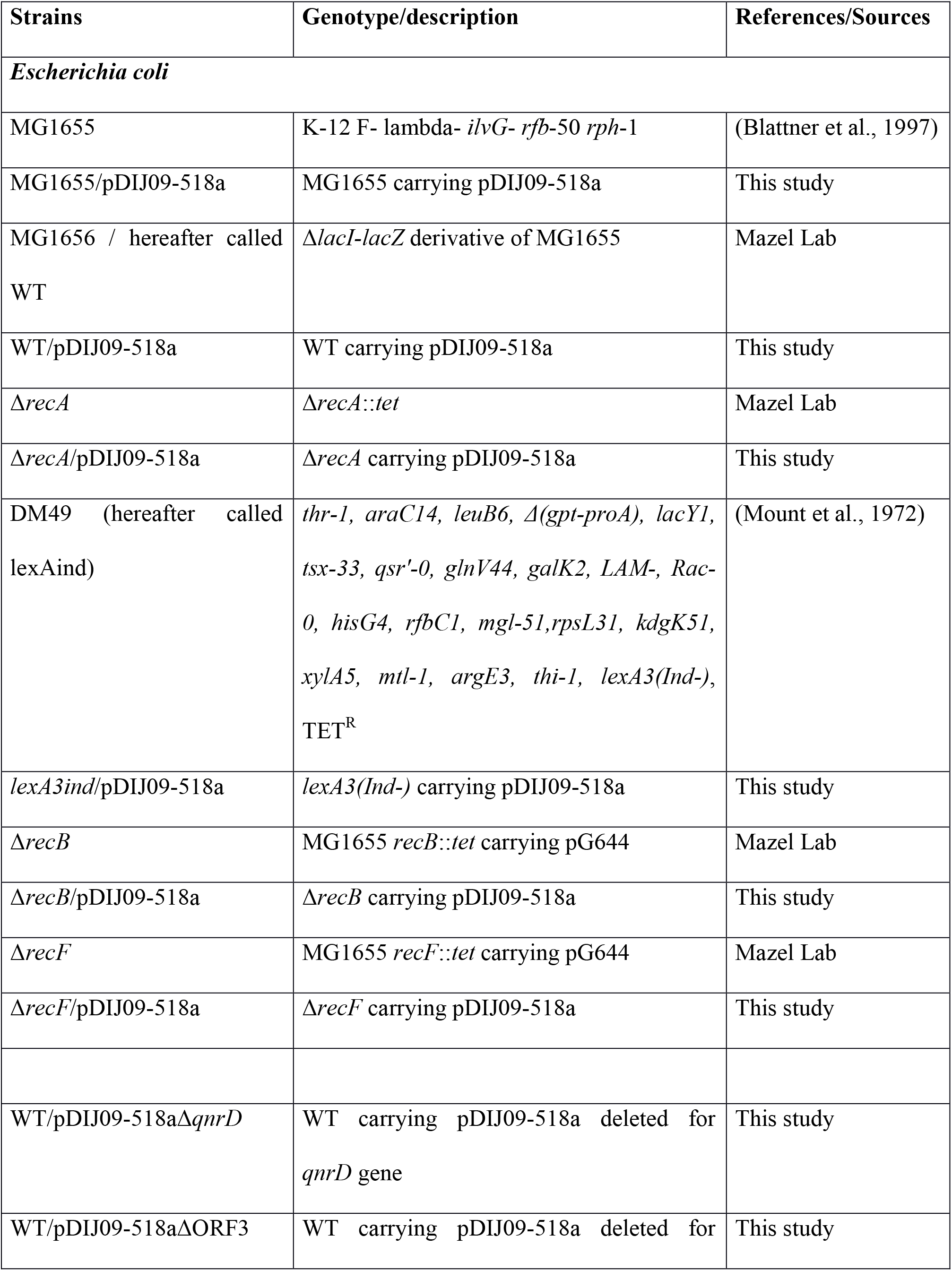
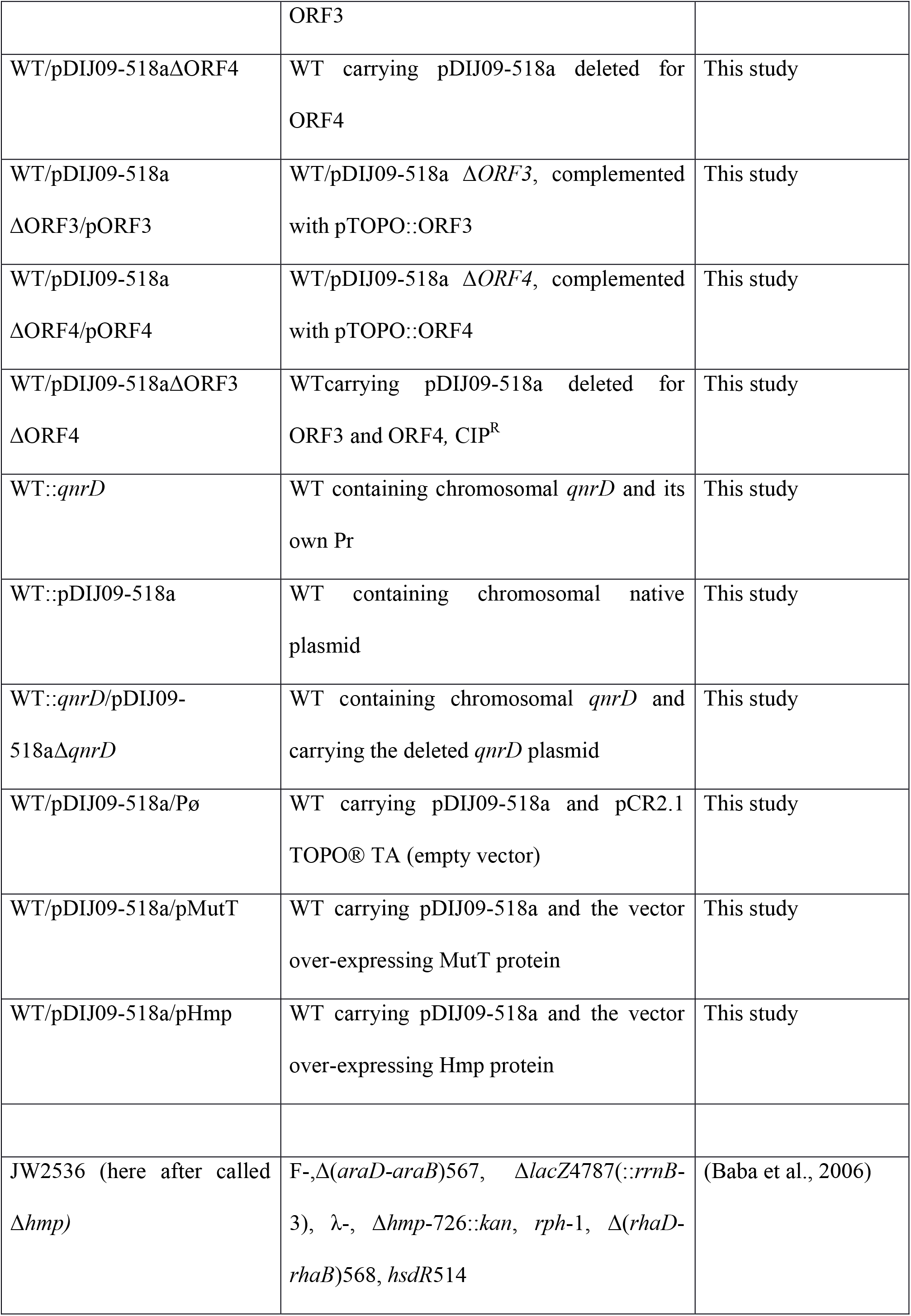
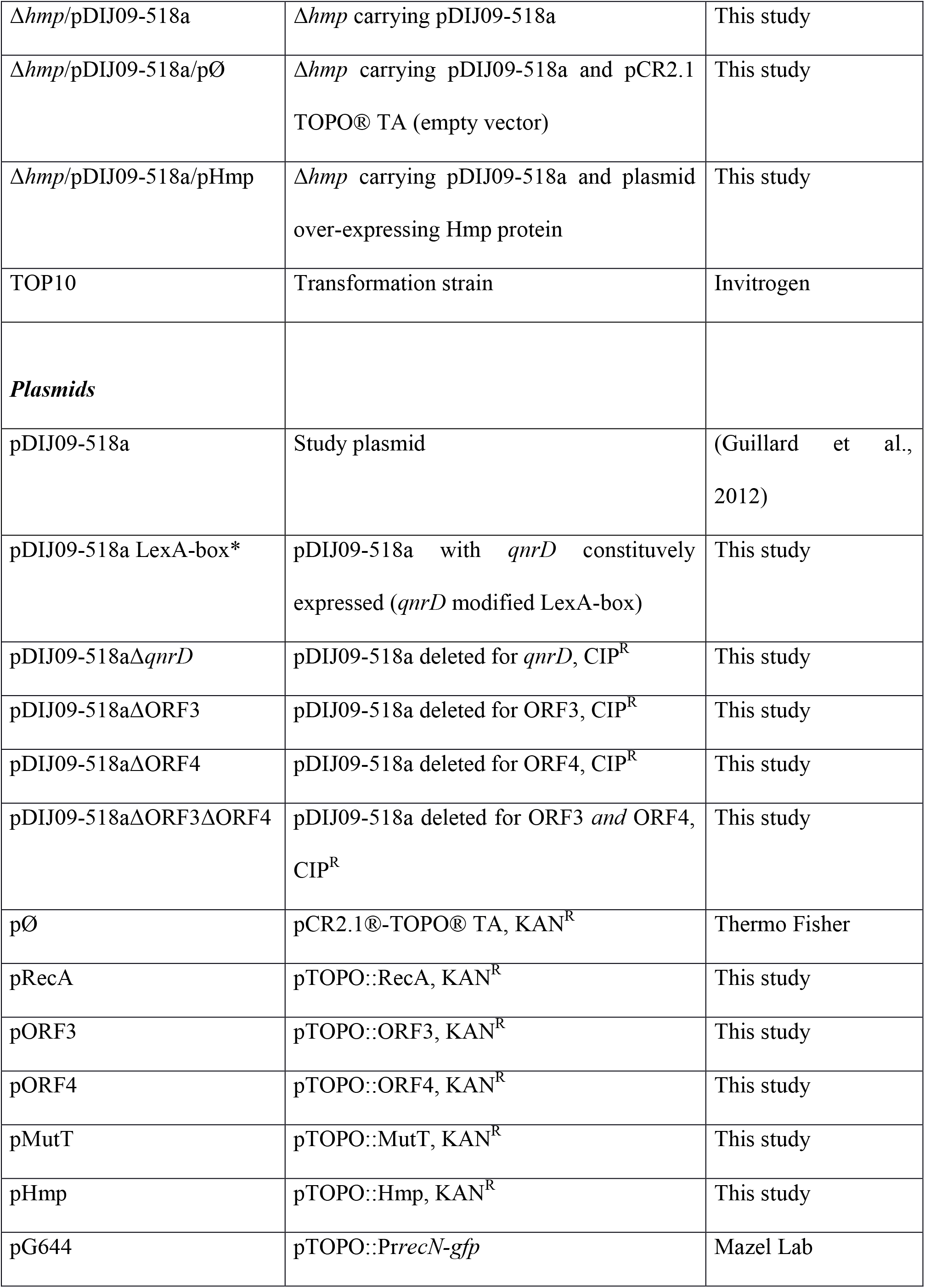
Strains and plasmids.

**Table S3.**
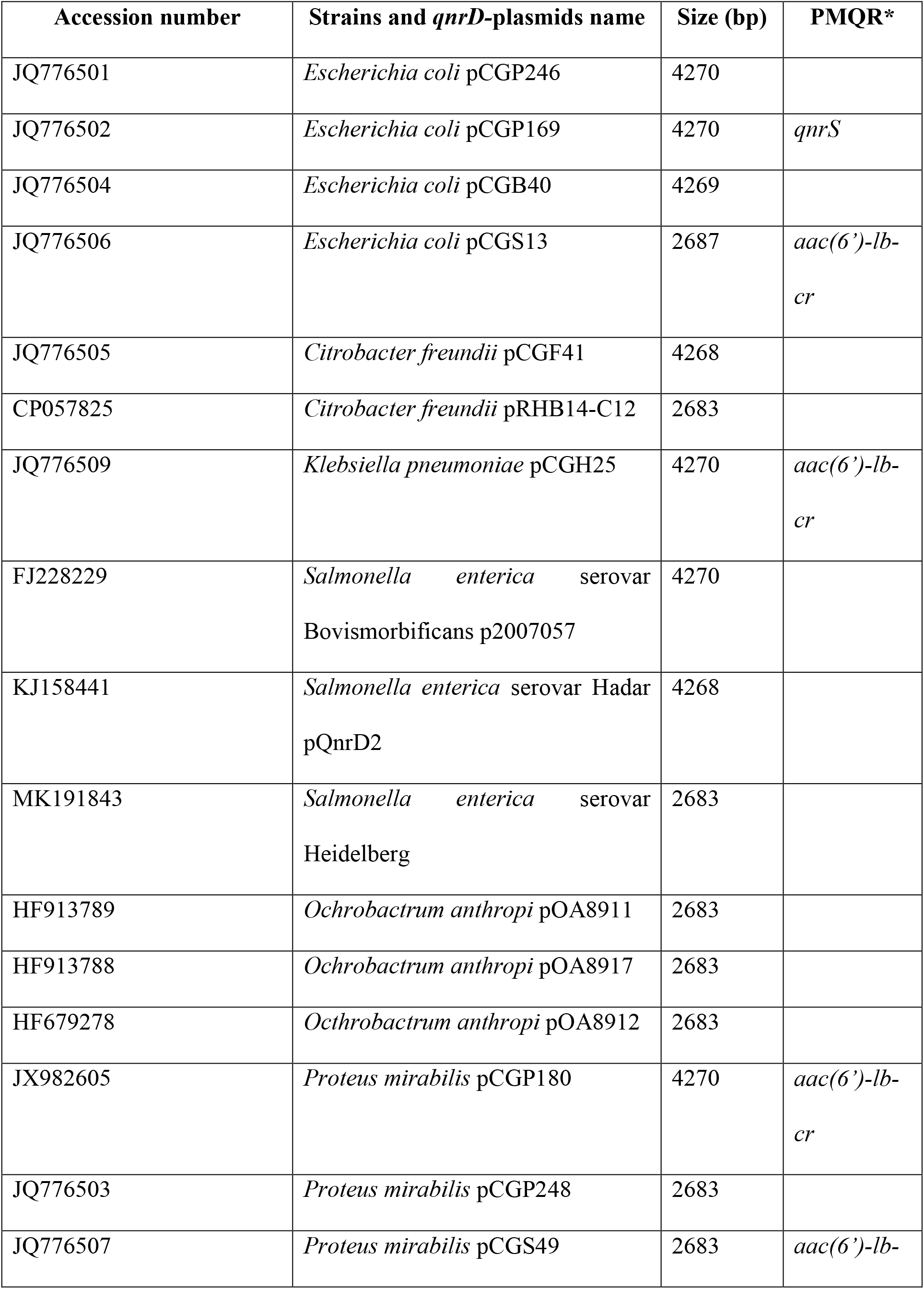
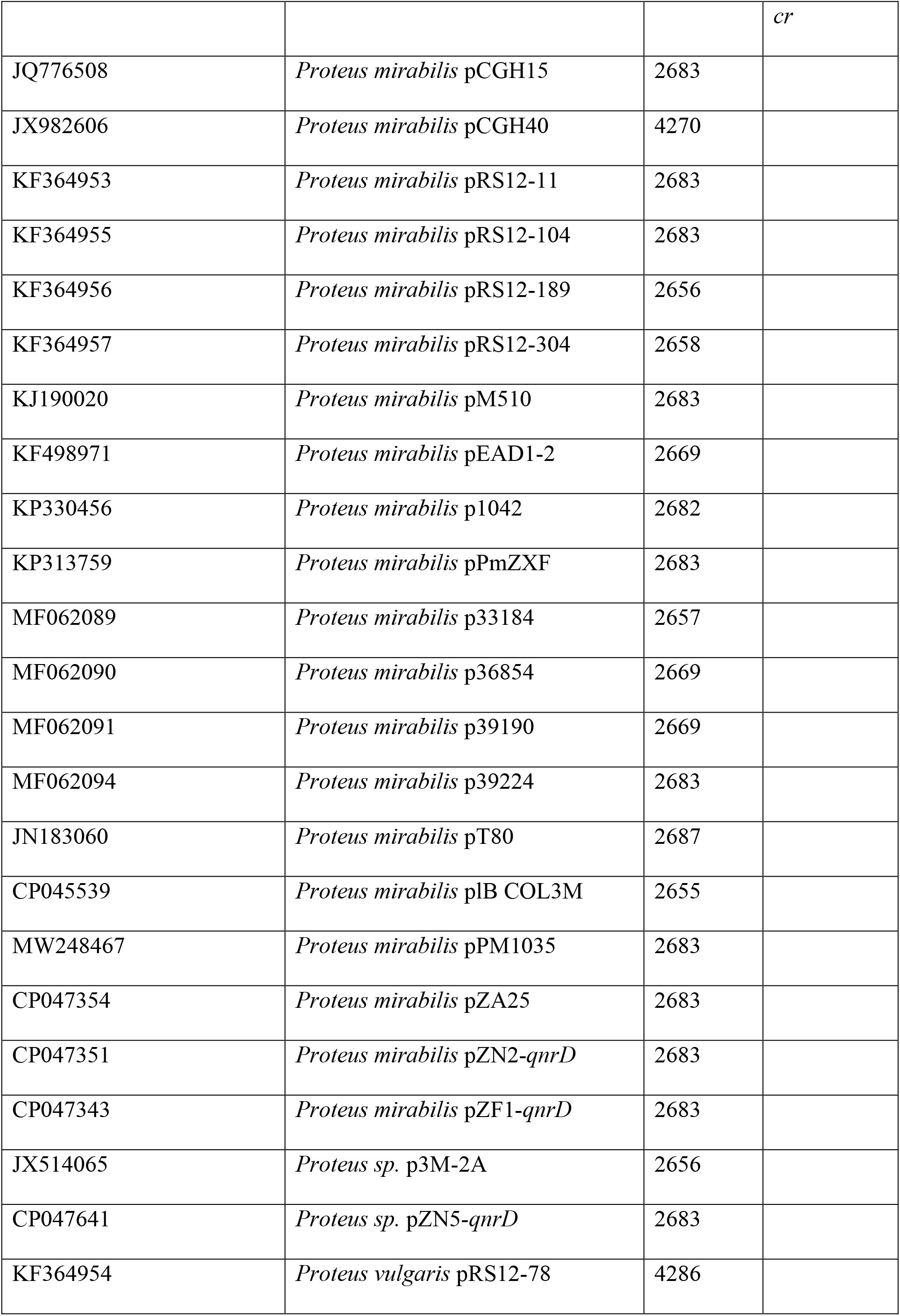
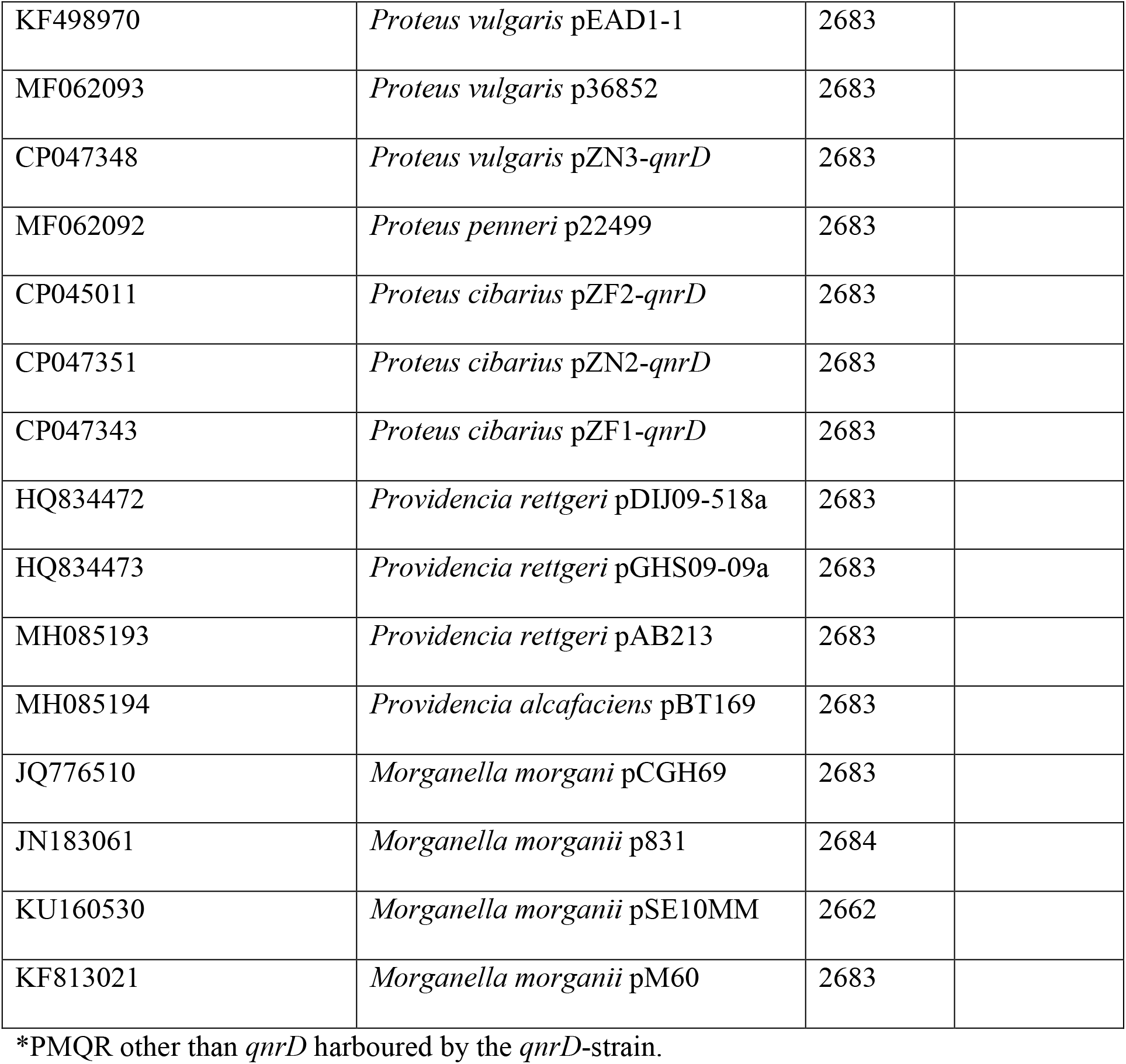
Genetic background of *qnrD* positive enterobacterial isolates.

| Accession number | Strains and *qnrD*-plasmids name | Size (bp) | PMQR* |
| --- | --- | --- | --- |
| JQ776501 | *Escherichia coli* pCGP246 | 4270 |  |
| JQ776502 | *Escherichia coli* pCGP169 | 4270 | *qnrS* |
| JQ776504 | *Escherichia coli* pCGB40 | 4269 |  |
| JQ776506 | *Escherichia coli* pCGS13 | 2687 | *aac(6')-lb-cr* |
| JQ776505 | *Citrobacter freundii* pCGF41 | 4268 |  |
| CP057825 | *Citrobacter freundii* pRHB14-C12 | 2683 |  |
| JQ776509 | *Klebsiella pneumoniae* pCGH25 | 4270 | *aac(6')-lb-cr* |
| FJ228229 | *Salmonella enterica* serovar<br>Bovismorbificans p2007057 | 4270 |  |
| KJ158441 | *Salmonella enterica* serovar Hadar<br>pQnrD2 | 4268 |  |
| MK191843 | *Salmonella enterica* serovar<br>Heidelberg | 2683 |  |
| HF913789 | *Ochrobactrum anthropi* pOA8911 | 2683 |  |
| HF913788 | *Ochrobactrum anthropi* pOA8917 | 2683 |  |
| HF679278 | *Ochrobactrum anthropi* pOA8912 | 2683 |  |
| JX982605 | *Proteus mirabilis* pCGP180 | 4270 | *aac(6')-lb-cr* |
| JQ776503 | *Proteus mirabilis* pCGP248 | 2683 |  |
| JQ776507 | *Proteus mirabilis* pCGS49 | 2683 | *aac(6')-lb-* *cr* |
| JQ776508 | *Proteus mirabilis* pCGH15 | 2683 |  |
| JX982606 | *Proteus mirabilis* pCGH40 | 4270 |  |
| KF364953 | *Proteus mirabilis* pRS12-11 | 2683 |  |
| KF364955 | *Proteus mirabilis* pRS12-104 | 2683 |  |
| KF364956 | *Proteus mirabilis* pRS12-189 | 2656 |  |
| KF364957 | *Proteus mirabilis* pRS12-304 | 2658 |  |
| KJ190020 | *Proteus mirabilis* pM510 | 2683 |  |
| KF498971 | *Proteus mirabilis* pEAD1-2 | 2669 |  |
| KP330456 | *Proteus mirabilis* p1042 | 2682 |  |
| KP313759 | *Proteus mirabilis* pPmZXF | 2683 |  |
| MF062089 | *Proteus mirabilis* p33184 | 2657 |  |
| MF062090 | *Proteus mirabilis* p36854 | 2669 |  |
| MF062091 | *Proteus mirabilis* p39190 | 2669 |  |
| MF062094 | *Proteus mirabilis* p39224 | 2683 |  |
| JN183060 | *Proteus mirabilis* pT80 | 2687 |  |
| CP045539 | *Proteus mirabilis* plB COL3M | 2655 |  |
| MW248467 | *Proteus mirabilis* pPM1035 | 2683 |  |
| CP047354 | *Proteus mirabilis* pZA25 | 2683 |  |
| CP047351 | *Proteus mirabilis* pZN2-*qnrD* | 2683 |  |
| CP047343 | *Proteus mirabilis* pZF1-*qnrD* | 2683 |  |
| JX514065 | *Proteus* sp. p3M-2A | 2656 |  |
| CP047641 | *Proteus* sp. pZN5-*qnrD* | 2683 |  |
| KF364954 | *Proteus vulgaris* pRS12-78 | 4286 |  |
| KF498970 | *Proteus vulgaris* pEAD1-1 | 2683 |  |
| MF062093 | *Proteus vulgaris* p36852 | 2683 |  |
| CP047348 | *Proteus vulgaris* pZN3-*qnrD* | 2683 |  |
| MF062092 | *Proteus penneri* p22499 | 2683 |  |
| CP045011 | *Proteus cibarius* pZF2-*qnrD* | 2683 |  |
| CP047351 | *Proteus cibarius* pZN2-*qnrD* | 2683 |  |
| CP047343 | *Proteus cibarius* pZF1-*qnrD* | 2683 |  |
| HQ834472 | *Providencia rettgeri* pDIJ09-518a | 2683 |  |
| HQ834473 | *Providencia rettgeri* pGHS09-09a | 2683 |  |
| MH085193 | *Providencia rettgeri* pAB213 | 2683 |  |
| MH085194 | *Providencia alcafaciens* pBT169 | 2683 |  |
| JQ776510 | *Morganella morganii* pCGH69 | 2683 |  |
| JN183061 | *Morganella morganii* p831 | 2684 |  |
| KU160530 | *Morganella morganii* pSE10MM | 2662 |  |
| KF813021 | *Morganella morganii* pM60 | 2683 |  |

\*PMQR other than *qnrD* harboured by the *qnrD*-strain.

**Table S4.**
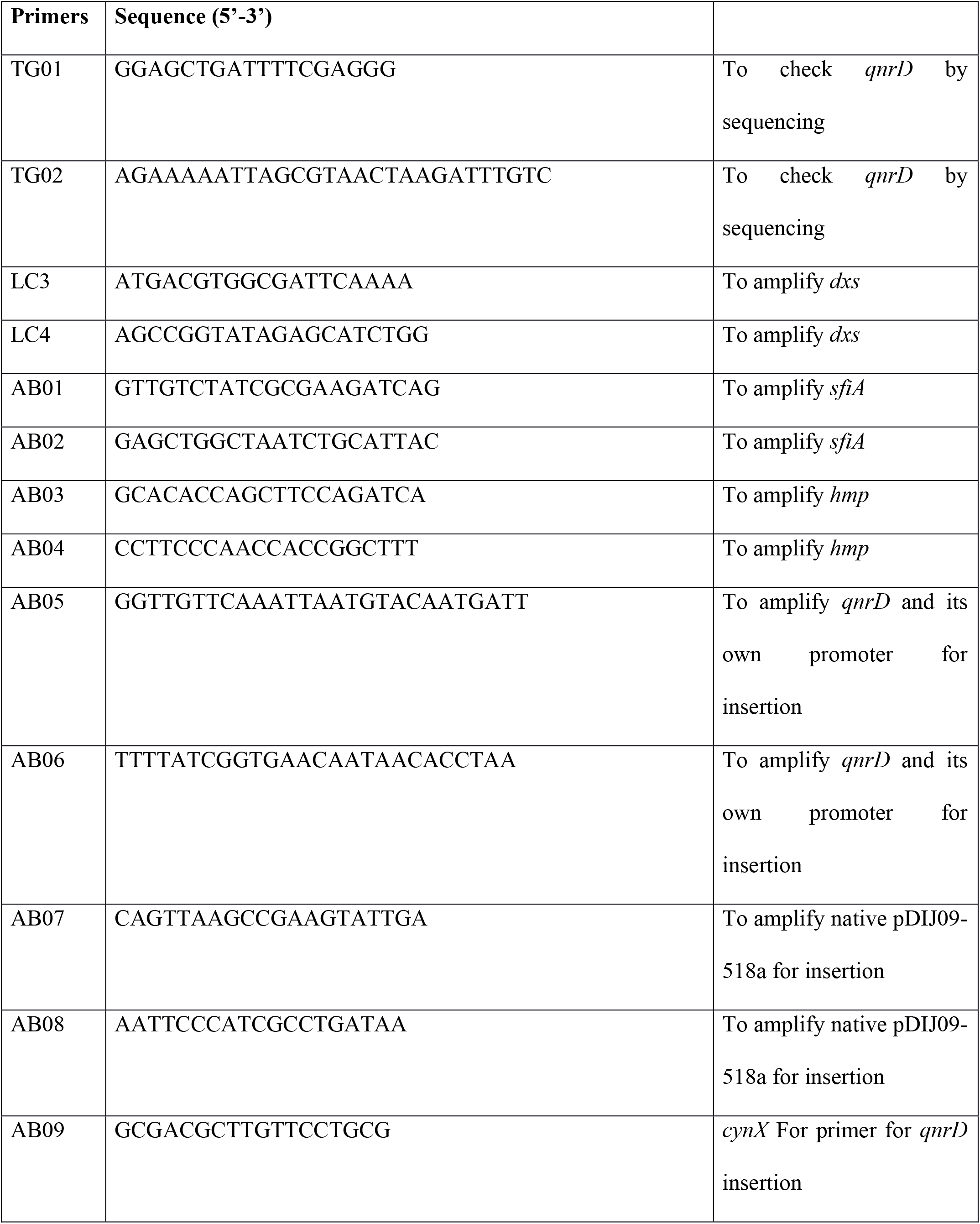
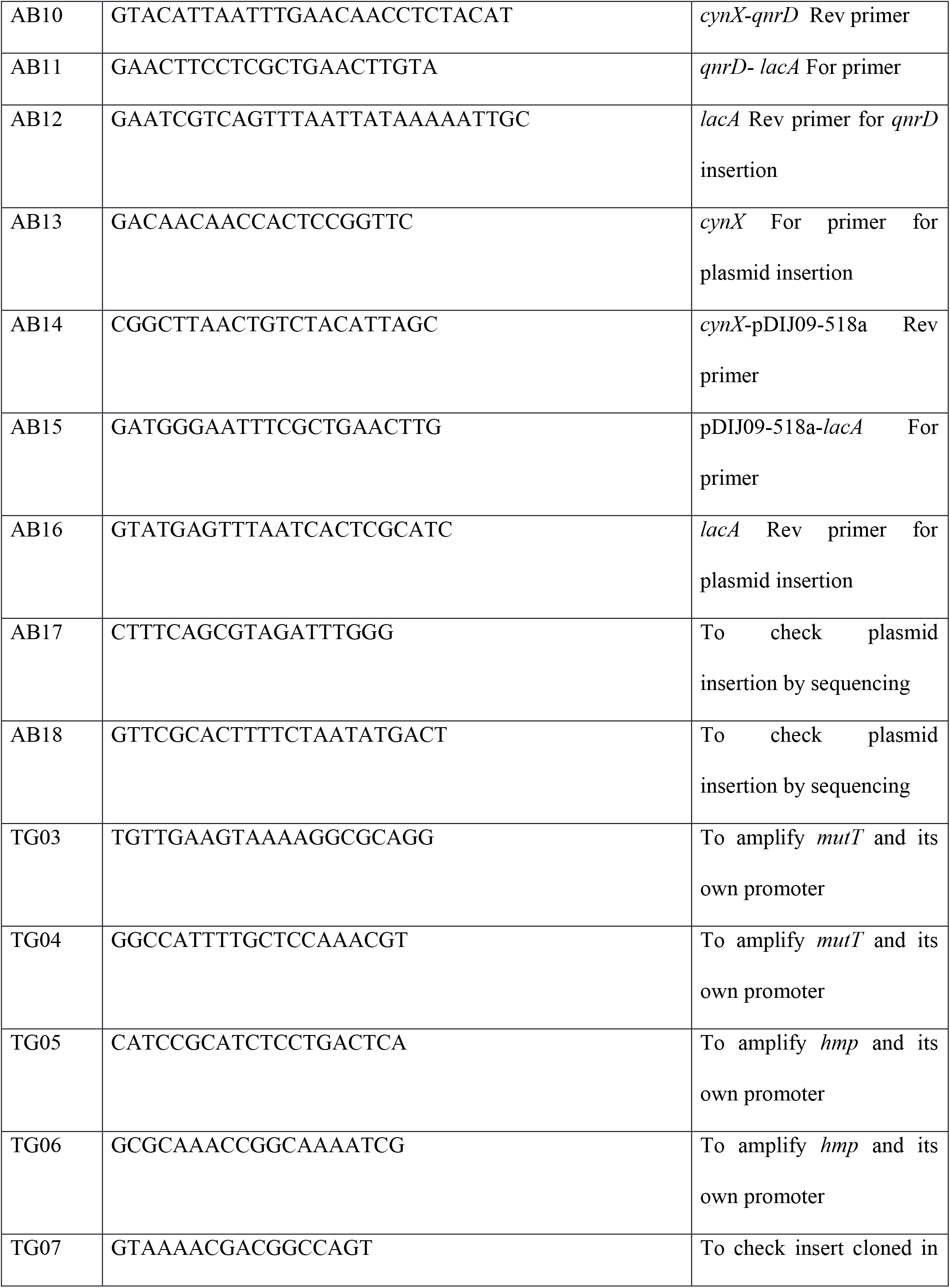
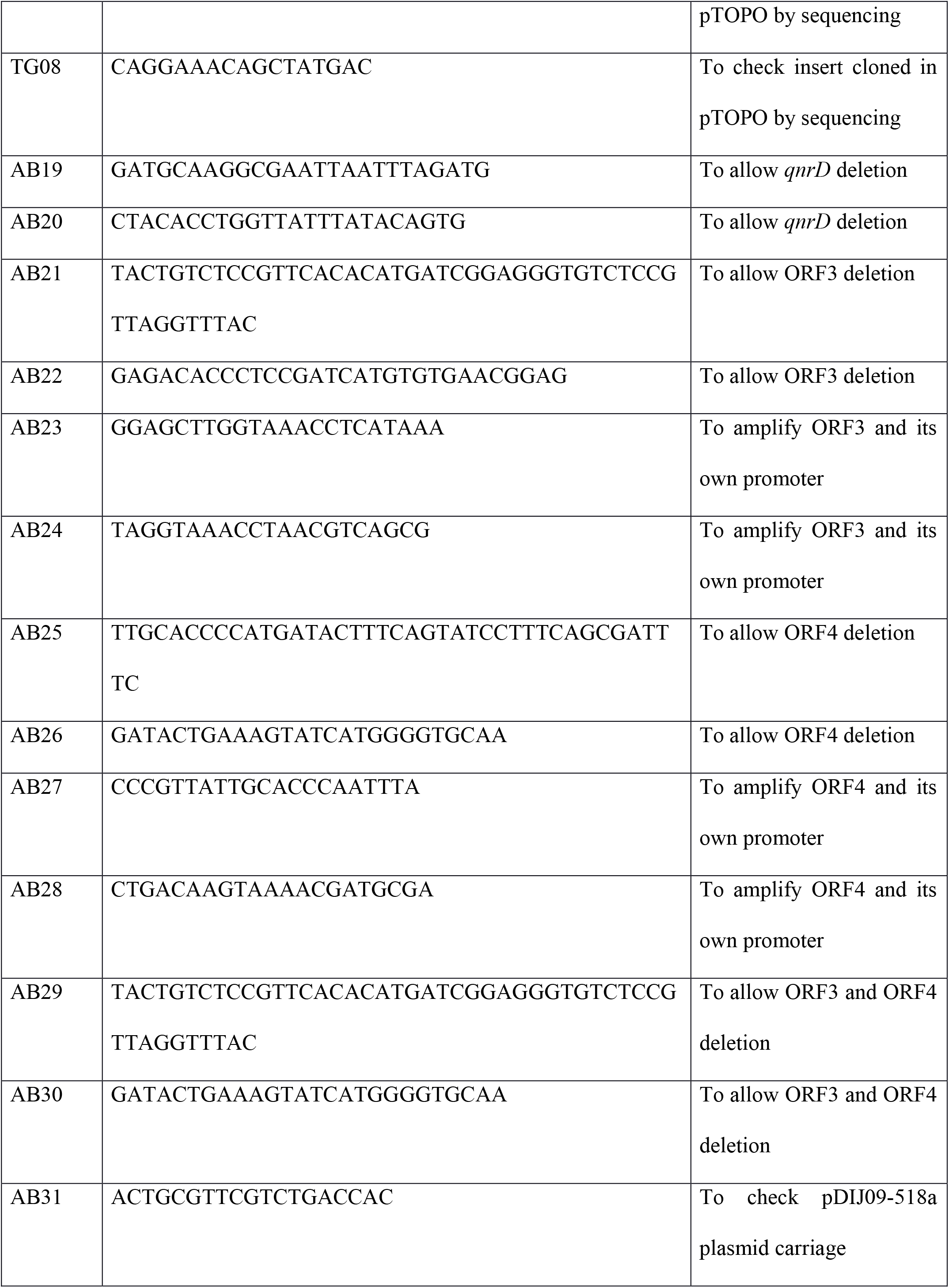
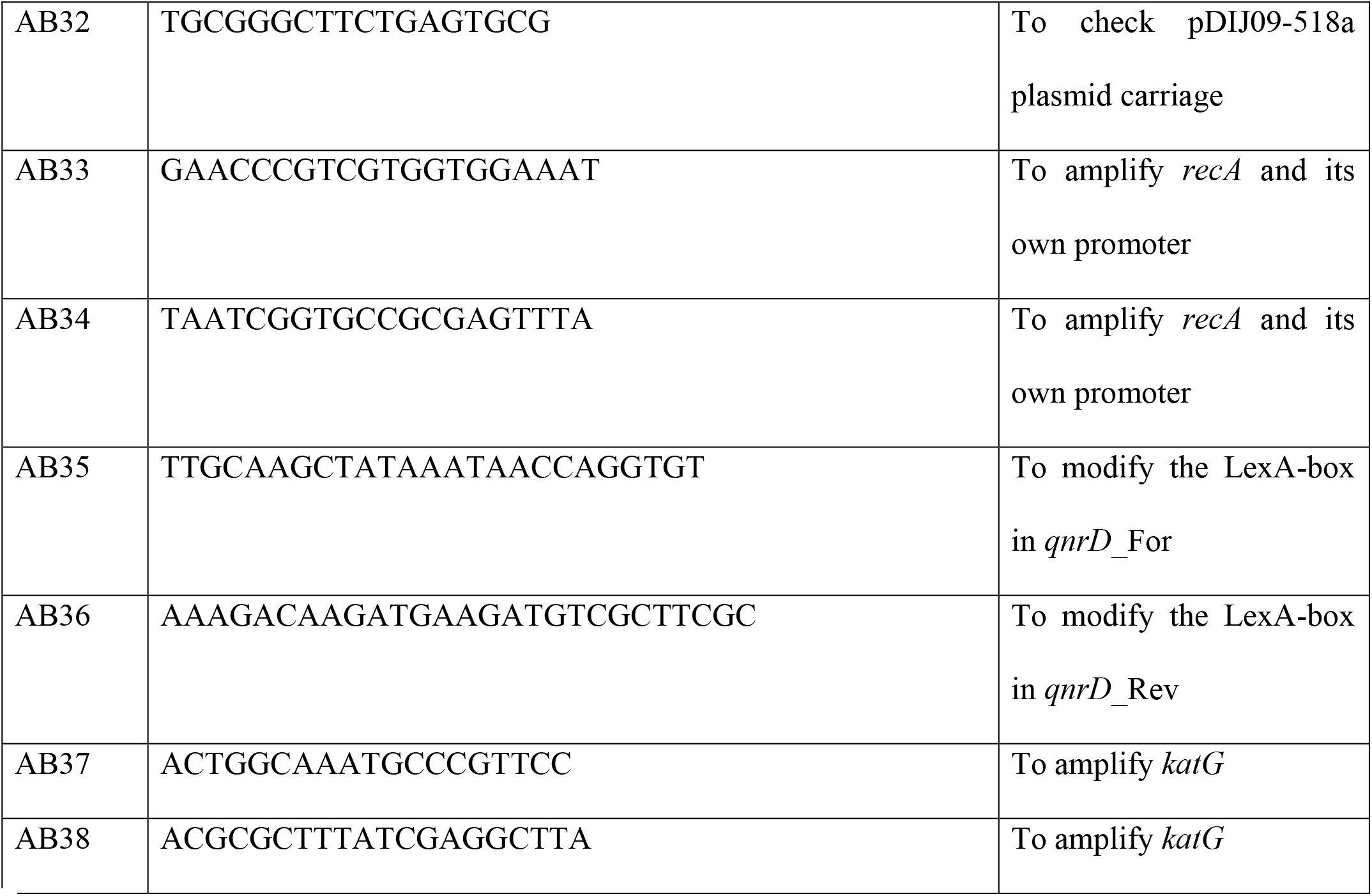
Primers used for this study.

**Fig. S1.**
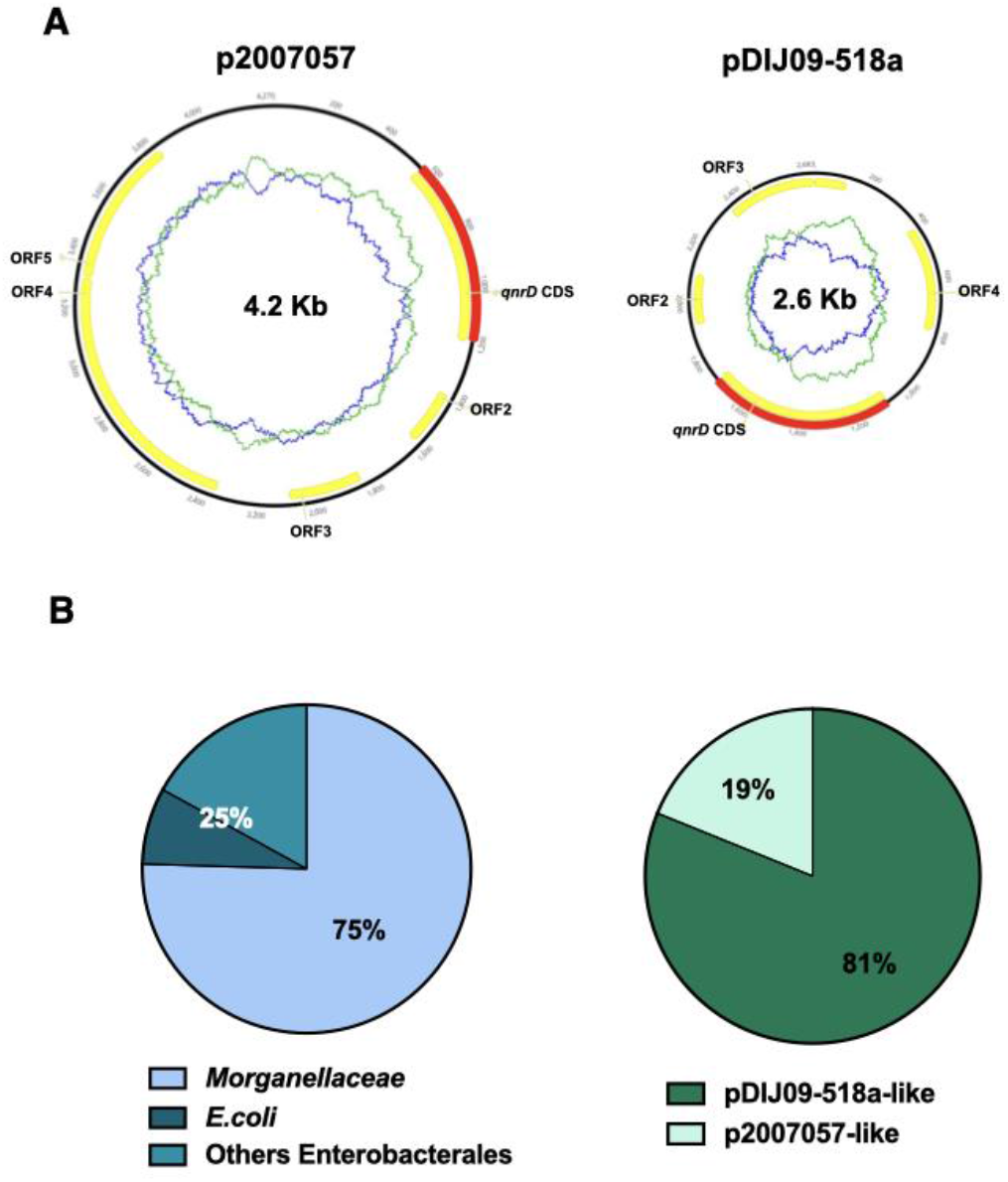
*qnrD* genes are carried by small plasmids. **(A)** The two *qnrD*-plasmid archetypes: p2007057 and pDIJ09-518a. **(B)** Distribution of the *qnrD*-plasmids among the 53 *qnrD* fully sequenced plasmids available in GenBank.

**Fig. S2.**
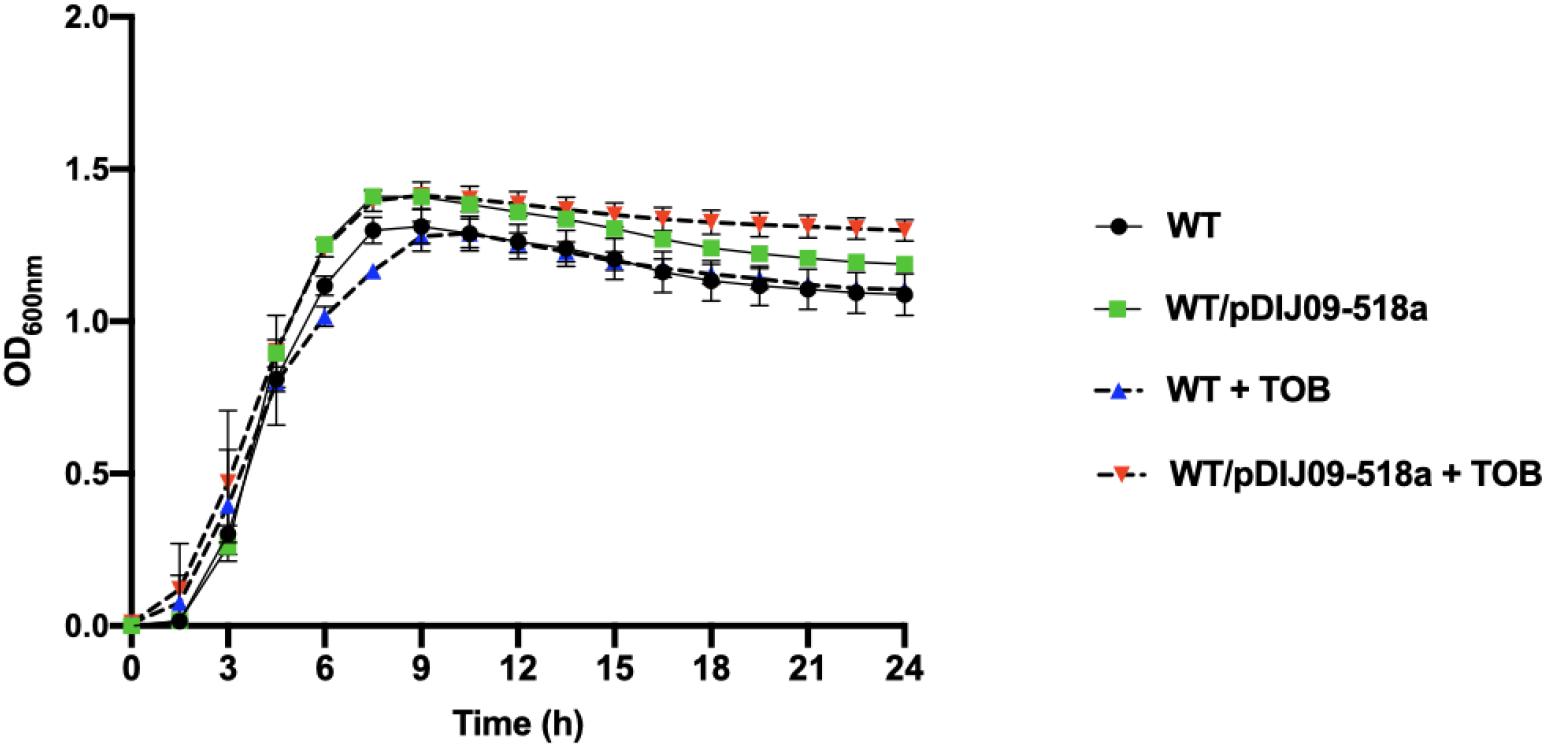
The viability of *E. coli* MG1656 carrying the *qnrD*-plasmid exposed to sub-MIC of tobramycin is not impaired and the plasmid is stable in an antibiotic free medium. The curves represent the OD at 600nm measured for 24h at 37°C, with shaking. The Y axis shows viability in the presence or in the absence of sub-MIC of tobramycin. Error bars represent standard deviation. Each strain was tested three times in triplicates.

**Fig. S3.**
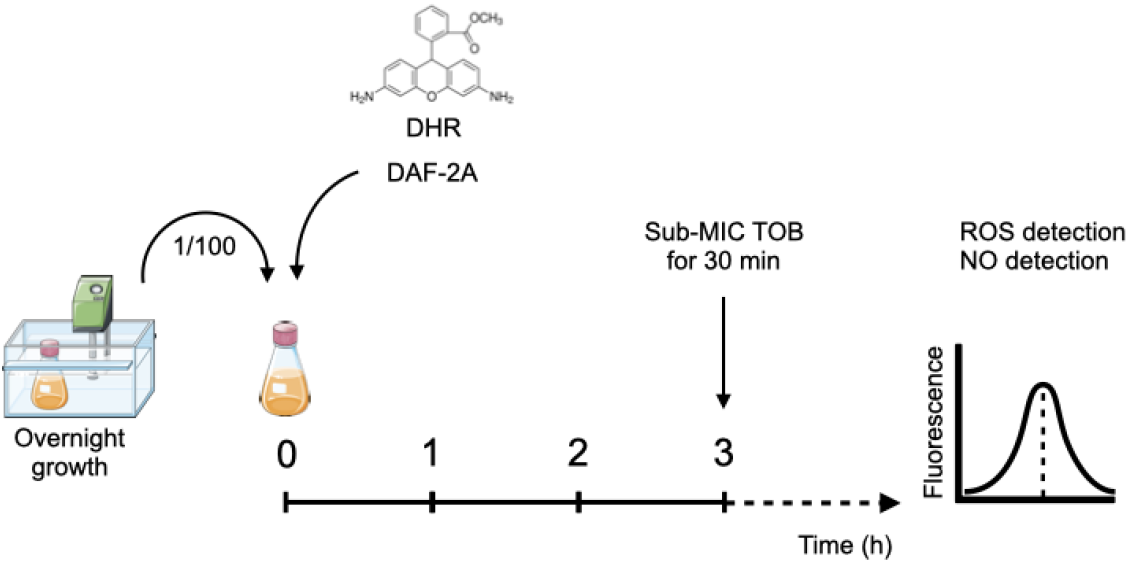
Schematic approach for fluorometric detection of free intercellular ROS and NO.

**Fig. S4.**
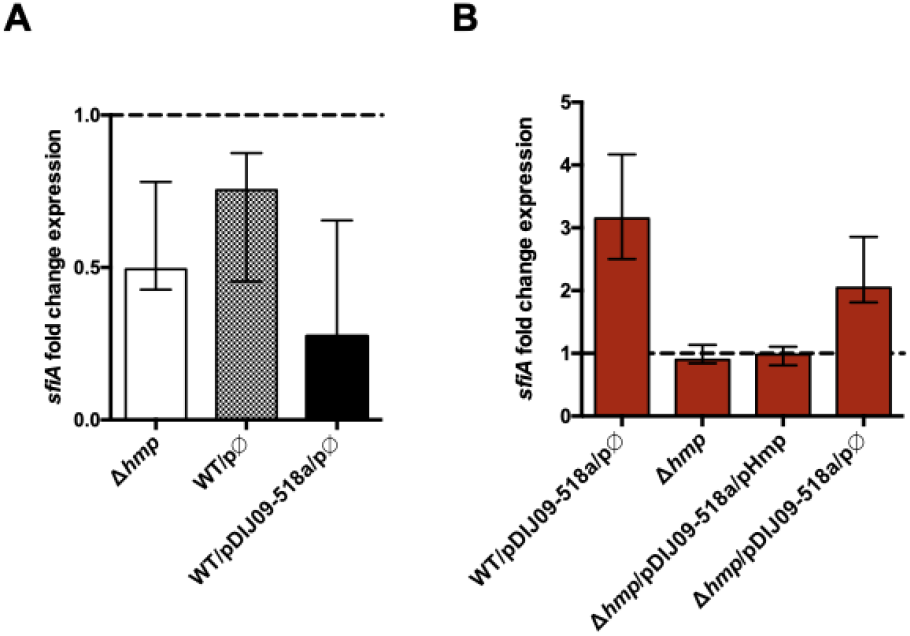
*hmp* deletion and empty vector carriage do not promote the SOS response induction. **(A)** Relative expression of *sfiA* in *hmp* deleted *E. coli* MG1656*, E. coli* MG1656 carrying the empty vector, in comparison to expression in wild-type (WT) *E. coli* MG1656 strain and *E. coli* co-carrying pDIJ09-518a and the empty vector compared to *E. coli* WT harbouring pDIJ09-518a, normalized with *dxs.* **(B)** Relative expression of *sfiA* in *hmp* mutant and derivatives strains exposed to sub-MIC concentration of tobramycin, in comparison to expression in LB, normalized with *dxs*. Data represent median values of 6 independent biological replicates, and error bars indicate upper/lower values. Source data are provided as a Source Data file.

